# Novel susceptibility loci and genetic regulation mechanisms for type 2 diabetes

**DOI:** 10.1101/284570

**Authors:** Angli Xue, Yang Wu, Zhihong Zhu, Futao Zhang, Kathryn E Kemper, Zhili Zheng, Loic Yengo, Luke R. Lloyd-Jones, Julia Sidorenko, Yeda Wu, eQTLGen Consortium, Allan F McRae, Peter M Visscher, Jian Zeng, Jian Yang

## Abstract

We conducted a meta-analysis of genome-wide association studies (GWAS) with ∼16 million genotyped/imputed genetic variants in 62,892 type 2 diabetes (T2D) cases and 596,424 controls of European ancestry. We identified 139 common and 4 rare (minor allele frequency < 0.01) variants associated with T2D, 42 of which (39 common and 3 rare variants) were independent of the known variants. Integration of the gene expression data from blood (*n* = 14,115 and 2,765) and other T2D-relevant tissues (*n* = up to 385) with the GWAS results identified 33 putative functional genes for T2D, three of which were targeted by approved drugs. A further integration of DNA methylation (*n* = 1,980) and epigenomic annotations data highlighted three putative T2D genes (*CAMK1D, TP53INP1* and *ATP5G1*) with plausible regulatory mechanisms whereby a genetic variant exerts an effect on T2D through epigenetic regulation of gene expression. We further found evidence that the T2D-associated loci have been under purifying selection.

## Introduction

Type 2 diabetes (T2D) is a common disease with a worldwide prevalence that increased rapidly from 4.7% in 1980 to 8.5% in 2014^1^. It is primarily caused by insulin resistance (failure of the body’s normal response to insulin) and/or insufficient insulin production by beta cells^2^. Genetic studies using linkage analysis and candidate gene approaches have led to the discovery of an initial set of T2D-associated loci (e.g., *PPARG, KCNJ11* and *TCF7L2*)^3-5^. Over the past decade, genome-wide association studies (GWAS) with increasing sample sizes have identified 144 genetic variants (not completely independent) at 129 loci associated with T2D^6-8^.

Despite the large number of variants discovered using GWAS, the associated variants in total explain only a small proportion (∼10%) of the heritability of T2D^9,10^. This well-known “missing heritability” problem is likely due to the presence of common variants (minor allele frequencies (MAF) > 0.01) that have small effects that have not yet been detected and/or rare variants that are not well tagged by common SNPs^9^. The contribution of rare variants to genetic variation in the occurrence of common diseases is under debate^11^, and a recent study suggested that the contribution of rare variants to the heritability of T2D is likely to be limited^12^. If most T2D- associated genetic variants are common in the population, continual discoveries of variants with small effects are expected from large-scale GWAS using the current experimental design. Furthermore, limited progress has been made in understanding the regulatory mechanisms of the genetic loci identified by GWAS. Thus, the etiology and the genetic basis underlying the development of the disease remain largely unknown. Recent methodological advances have provided us with an opportunity to identify functional genes and their regulatory elements by combining GWAS summary statistics with data from molecular quantitative trait loci studies with large sample size^13-15^.

In this study, we performed a meta-analysis of GWAS with the largest sample size for T2D to date (62,892 cases and 596,424 controls), by combining three large GWAS data sets: DIAbetes Genetics Replication And Meta-analysis (DIAGRAM)^7^, Genetic Epidemiology Research on Aging (GERA)^16^ and the full cohort release of the UK Biobank (UKB)^17^. We then integrated the GWAS meta-analysis results with gene expression and DNA methylation data to identify genes that might be functionally relevant to T2D and to infer plausible mechanisms whereby genetic variants affect T2D risk through gene regulation by DNA methylation^15^. We further estimated the genetic architecture of T2D using whole-genome estimation approaches.

## Results

### Meta-analysis identifies 39 previously unknown loci

We meta-analyzed 5,053,015 genotyped or imputed autosomal SNPs (MAF > 0.01) in 62,892 T2D cases and 596,424 controls from the DIAGRAM (12,171 cases vs. 56,862 controls in stage 1 and 22,669 cases vs. 58,119 controls in stage 2), GERA (6,905 cases and 46,983 controls) and UKB (21,147 cases and 434,460 controls) data sets after quality controls (**Supplementary Fig. 1** and **Methods**). Summary statistics in DIAGRAM were imputed to the 1000 Genomes Project^18^ (1KGP) phase 1 using a summary-data-based imputation approach, ImpG^19^ (**Supplementary Note 1**), and we used an inverse-variance method^20^ to meta-analyze the imputed DIAGRAM data with the summary data from GWAS analyses of GERA (1KGP imputed data) and UKB (Haplotype Reference Consortium^21^ or HRC imputed data) (**Methods** and **Fig. 1a**). All the individuals except for a Pakistani cohort in DIAGRAM stage 2 (see **Methods**) were of European ancestry. We demonstrated by linkage disequilibrium (LD) score regression analysis^22,23^ that the inflation in test statistics due to population structure was negligible in each data set, and there was no evidence of sample overlap among the three data sets (**Supplementary Note 2** and **Supplementary Table 1**). The mean χ^2^ statistic was 1.685. LD score regression analysis of the meta-analysis summary statistics showed an estimate of SNP-based heritability 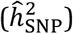 on the liability scale of 0.196 (*s.e.* = 0.011) and an estimate of intercept of 1.049 (*s.e.* = 0.014), consistent with a model in which the genomic inflation in test statistics is driven by polygenic effects^22,24^. After clumping the SNPs using LD information from the UKB genotypes (clumping *r*^2^ threshold = 0.01 and window size = 1 Mb), there were 139 near-independent variants at *P* < 5×10^−8^ (**Supplementary Table 2**). All of the loci previously reported by DIAGRAM were still genome-wide significant in our meta-analysis results. The most significant association was at rs7903146 (*P* = 1.3×10^−347^) at the known *TCF7L2* locus^5,25^. Among the 139 variants, 39 are not in LD with the known variants (**Fig. 1** and **Table 1**). The result remained unchanged when the GERA cohort was imputed to HRC (**Supplementary Fig. 2**). We regarded these 39 variants as novel discoveries; more than half of them passed a more stringent significance threshold at *P* < 1×10^−8^ (**Table 1**), a conservative control of genome-wide false positive rate (GWFPR) suggested by a recent simulation study^26^. The functional relevance of some novel gene loci to the disease is supported by existing biological or molecular evidence related to insulin and glucose (**Supplementary Note 3**). Forest plots showed that the effect directions of the 39 novel loci were consistent across the three GWAS data sets (**Supplementary Fig. 3**). Regional association plots show that some loci have complicated LD structures, and it is largely unclear which genes are responsible for the observed SNP-T2D associations (**Supplementary Fig. 4**). We also performed gene-based analysis by GCTA-fastBAT^27^ and conditional analysis by GCTA-COJO^28^ and discovered four loci with multiple independent signals associated with T2D (**Supplementary Note 4-5, Supplementary Fig. 5** and **Supplementary Tables 3-5**).

**Table 1.**
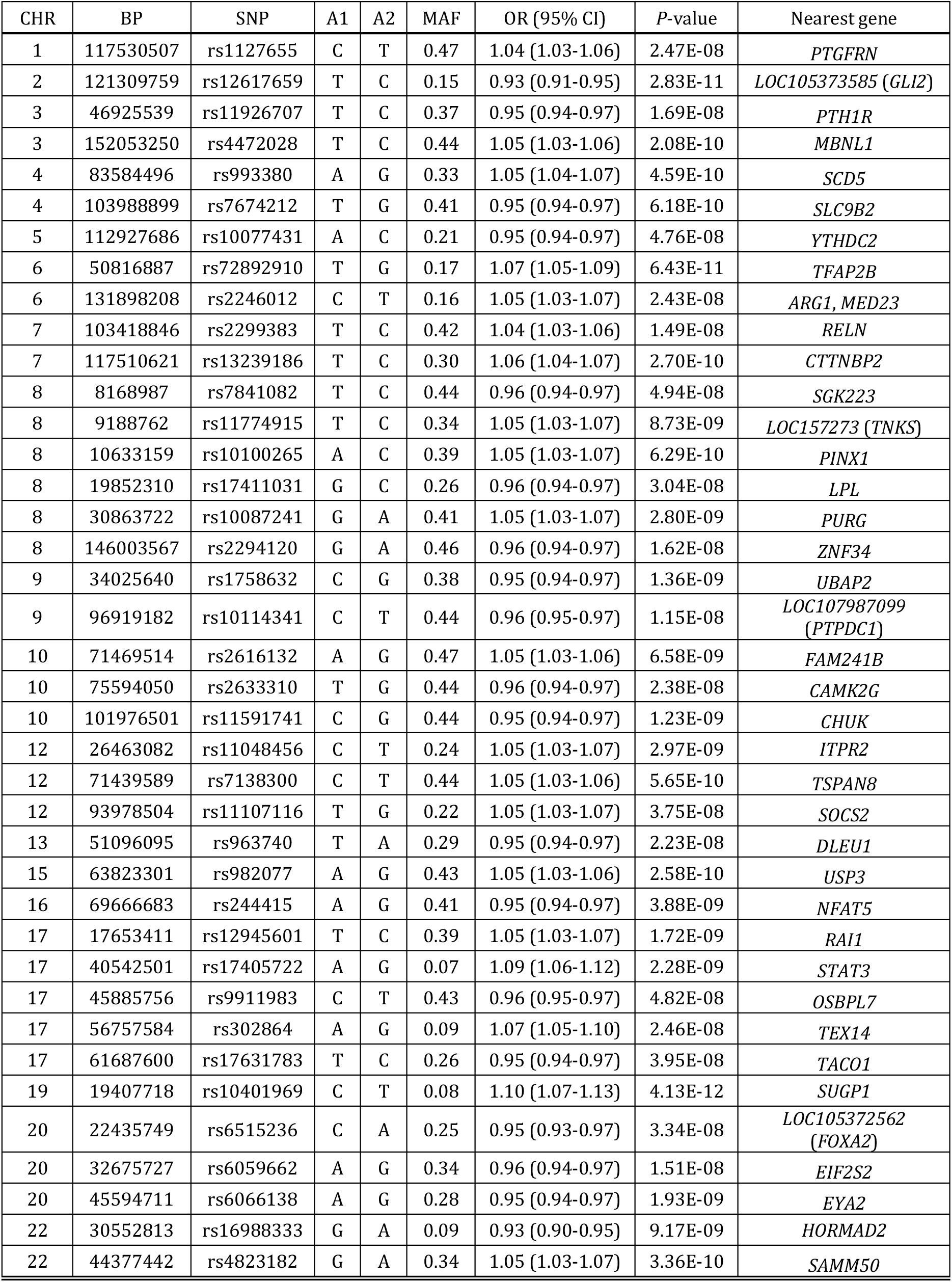
Common variants at 39 previously unknown T2D-associated loci

| CHR | BP | SNP | A1 | A2 | MAF | OR (95% CI) | P-value | Nearest gene |
| --- | --- | --- | --- | --- | --- | --- | --- | --- |
| 1 | 117530507 | rs1127655 | C | T | 0.47 | 1.04 (1.03-1.06) | 2.47E-08 | *PTGFRN* |
| 2 | 121309759 | rs12617659 | T | C | 0.15 | 0.93 (0.91-0.95) | 2.83E-11 | *LOC105373585 (GLI2)* |
| 3 | 46925539 | rs11926707 | T | C | 0.37 | 0.95 (0.94-0.97) | 1.69E-08 | *PTH1R* |
| 3 | 152053250 | rs4472028 | T | C | 0.44 | 1.05 (1.03-1.06) | 2.08E-10 | *MBNL1* |
| 4 | 83584496 | rs993380 | A | G | 0.33 | 1.05 (1.04-1.07) | 4.59E-10 | *SCD5* |
| 4 | 103988899 | rs7674212 | T | G | 0.41 | 0.95 (0.94-0.97) | 6.18E-10 | *SLC9B2* |
| 5 | 112927686 | rs10077431 | A | C | 0.21 | 0.95 (0.94-0.97) | 4.76E-08 | *YTHDC2* |
| 6 | 50816887 | rs72892910 | T | G | 0.17 | 1.07 (1.05-1.09) | 6.43E-11 | *TFAP2B* |
| 6 | 131898208 | rs2246012 | C | T | 0.16 | 1.05 (1.03-1.07) | 2.43E-08 | *ARG1, MED23* |
| 7 | 103418846 | rs2299383 | T | C | 0.42 | 1.04 (1.03-1.06) | 1.49E-08 | *RELN* |
| 7 | 117510621 | rs13239186 | T | C | 0.30 | 1.06 (1.04-1.07) | 2.70E-10 | *CTTNBP2* |
| 8 | 8168987 | rs7841082 | T | C | 0.44 | 0.96 (0.94-0.97) | 4.94E-08 | *SGK223* |
| 8 | 9188762 | rs11774915 | T | C | 0.34 | 1.05 (1.03-1.07) | 8.73E-09 | *LOC157273 (TNKS)* |
| 8 | 10633159 | rs10100265 | A | C | 0.39 | 1.05 (1.03-1.07) | 6.29E-10 | *PINX1* |
| 8 | 19852310 | rs17411031 | G | C | 0.26 | 0.96 (0.94-0.97) | 3.04E-08 | *LPL* |
| 8 | 30863722 | rs10087241 | G | A | 0.41 | 1.05 (1.03-1.07) | 2.80E-09 | *PURG* |
| 8 | 146003567 | rs2294120 | G | A | 0.46 | 0.96 (0.94-0.97) | 1.62E-08 | *ZNF34* |
| 9 | 34025640 | rs1758632 | C | G | 0.38 | 0.95 (0.94-0.97) | 1.36E-09 | *UBAP2* |
| 9 | 96919182 | rs10114341 | C | T | 0.44 | 0.96 (0.95-0.97) | 1.15E-08 | *LOC107987099 (PTPDC1)* |
| 10 | 71469514 | rs2616132 | A | G | 0.47 | 1.05 (1.03-1.06) | 6.58E-09 | *FAM241B* |
| 10 | 75594050 | rs2633310 | T | G | 0.44 | 0.96 (0.94-0.97) | 2.38E-08 | *CAMK2G* |
| 10 | 101976501 | rs11591741 | C | G | 0.44 | 0.95 (0.94-0.97) | 1.23E-09 | *CHUK* |
| 12 | 26463082 | rs11048456 | C | T | 0.24 | 1.05 (1.03-1.07) | 2.97E-09 | *ITPR2* |
| 12 | 71439589 | rs7138300 | C | T | 0.44 | 1.05 (1.03-1.06) | 5.65E-10 | *TSPAN8* |
| 12 | 93978504 | rs11107116 | T | G | 0.22 | 1.05 (1.03-1.07) | 3.75E-08 | *SOCS2* |
| 13 | 51096095 | rs963740 | T | A | 0.29 | 0.95 (0.94-0.97) | 2.23E-08 | *DLEU1* |
| 15 | 63823301 | rs982077 | A | G | 0.43 | 1.05 (1.03-1.06) | 2.58E-10 | *USP3* |
| 16 | 69666683 | rs244415 | A | G | 0.41 | 0.95 (0.94-0.97) | 3.88E-09 | *NFAT5* |
| 17 | 17653411 | rs12945601 | T | C | 0.39 | 1.05 (1.03-1.07) | 1.72E-09 | *RAI1* |
| 17 | 40542501 | rs17405722 | A | G | 0.07 | 1.09 (1.06-1.12) | 2.28E-09 | *STAT3* |
| 17 | 45885756 | rs9911983 | C | T | 0.43 | 0.96 (0.95-0.97) | 4.82E-08 | *OSBPL7* |
| 17 | 56757584 | rs302864 | A | G | 0.09 | 1.07 (1.05-1.10) | 2.46E-08 | *TEX14* |
| 17 | 61687600 | rs17631783 | T | C | 0.26 | 0.95 (0.94-0.97) | 3.95E-08 | *TACO1* |
| 19 | 19407718 | rs10401969 | C | T | 0.08 | 1.10 (1.07-1.13) | 4.13E-12 | *SUGP1* |
| 20 | 22435749 | rs6515236 | C | A | 0.25 | 0.95 (0.93-0.97) | 3.34E-08 | *LOC105372562 (FOXA2)* |
| 20 | 32675727 | rs6059662 | A | G | 0.34 | 0.96 (0.94-0.97) | 1.51E-08 | *EIF2S2* |
| 20 | 45594711 | rs6066138 | A | G | 0.28 | 0.95 (0.94-0.97) | 1.93E-09 | *EYA2* |
| 22 | 30552813 | rs16988333 | G | A | 0.09 | 0.93 (0.90-0.95) | 9.17E-09 | *HORMAD2* |
| 22 | 44377442 | rs4823182 | G | A | 0.34 | 1.05 (1.03-1.07) | 3.36E-10 | *SAMM50* |

**Table 2.**
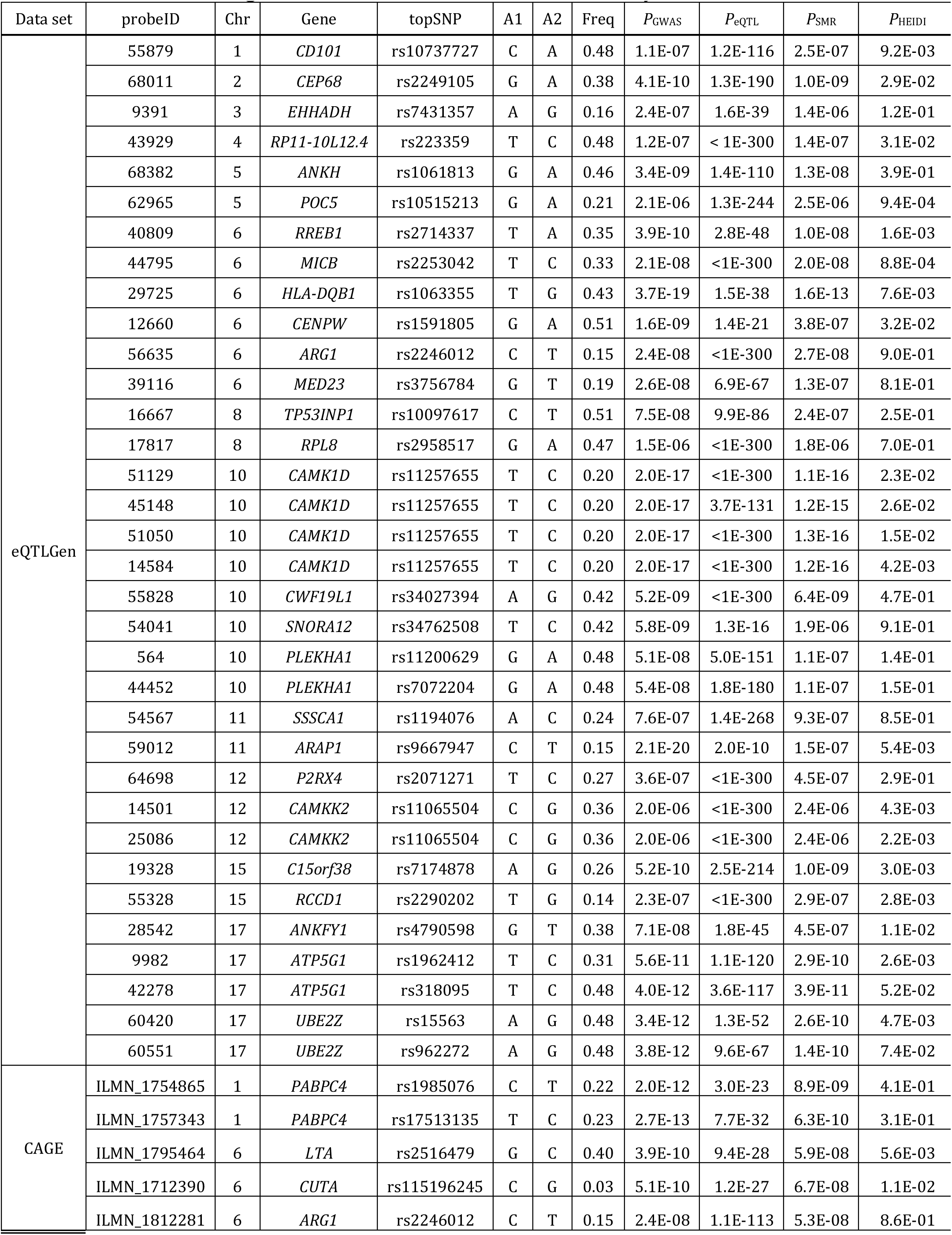
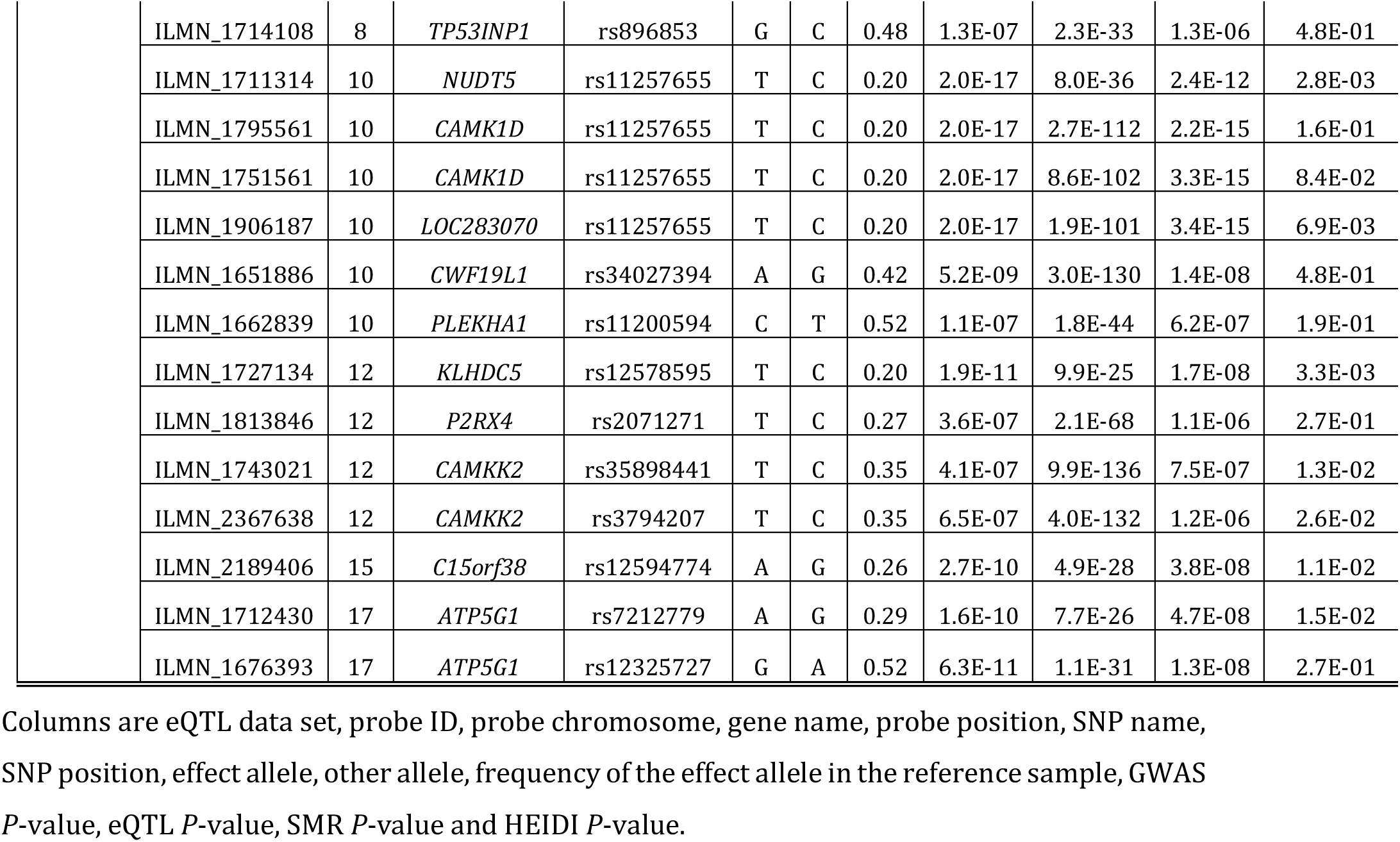
Putative functional genes for T2D identified from the SMR analysis

**Figure 1.**
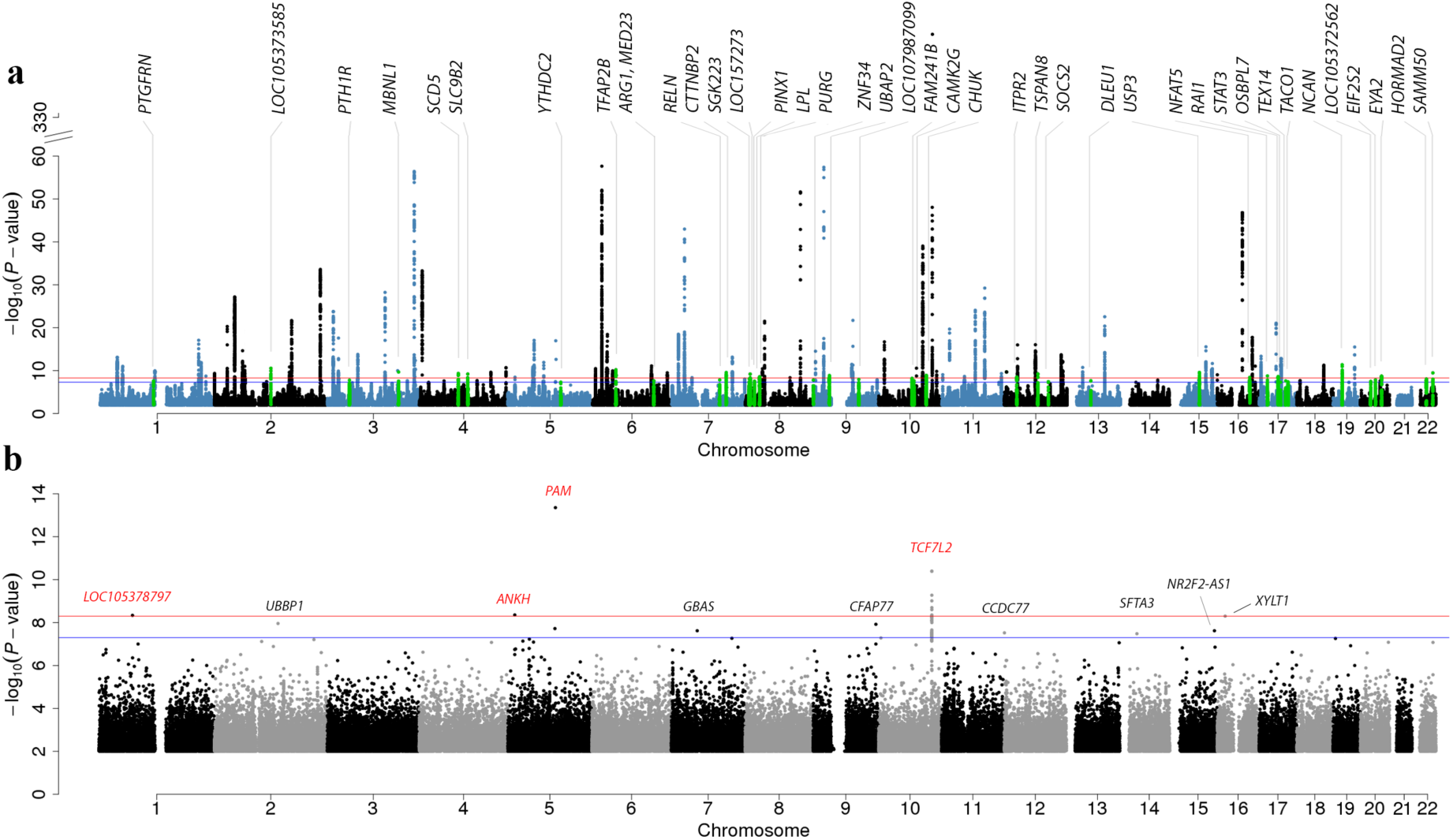
Manhattan plot of common variants identified by the meta-analysis and rare variants identified by a GWAS analysis in UKB. a) GWAS results for common variants (MAF > 0.01) in the meta-analysis. The 39 novel loci are annotated and highlighted in green. b) GWAS results of rare variants (0.0001 < MAF < 0.01) in UKB. Four loci with *P* < 5×10^−9^ are highlighted in red. For better graphical presentation, SNPs with 1×10^−60^ < *P*_meta_ < 1×10^−330^ and *P*_meta_ > 1×10^−2^ have been omitted from both panels. The blue lines denote the genome-wide significant threshold of *P* < 5×10^−8^, and the red lines denote a more stringent threshold of *P* < 5×10^−9^

Of all the 139 T2D-associated loci identified in our meta-analysis, 16 and 25 were significant in insulin secretion and sensitivity GWAS, respectively, from the MAGIC consortium^29,30^ (**URLs**) after correcting for multiple tests (*i.e.*, 0.05 / 139), with only one locus showing significant associations with both insulin secretion and sensitivity. The limited number of overlapping associations observed might be due to the relatively small sample sizes in the insulin studies. We further estimated the genetic correlation (*r*_g_) between insulin secretion (or sensitivity) and T2D by the bivariate LD score regression approach^23^ using summary-level data. The estimate of *r*_g_ between T2D and insulin secretion was −0.15 (*s.e.* = 0.10), and that between T2D and insulin sensitivity was −0.57 (*s.e.* = 0.10).

### Rare variants associated with T2D

Very few rare variants associated with T2D have been identified in previous studies^31-35^. We included 10,849,711 rare variants (0.0001 < MAF < 0.01) in the association analysis in UKB and detected 11 rare variants at *P* < 5×10^−8^ and 4 of them were at *P* < 5×10^−9^ (**Fig. 1b** and **Supplementary Table 6**). We focused only on the 4 signals at *P* < 5×10^−9^ because a recent study suggested that a *P*-value threshold of 5×10^−9^ is required to control a GWFPR at 0.05 in GWAS including both common and rare variants imputed from a fully sequenced reference^26^. Three of the rare variants were located at loci with significant common variant associations. The rs78408340 (OR = 1.33, *P* = 4.4×10^−14^) is a missense variant that encodes a p.Ser539Trp alteration in *PAM* and was reported to be associated with decreased insulin release from pancreatic beta cells^32^. Variant rs146886108 (odds ratio (OR) = 0.72, *P* = 4.4×10^−9^) is a novel locus, which showed a protective effect against T2D, is a missense variant that encodes p.Arg187Gln in *ANKH*^36^. Variant rs117229942 (OR = 0.70, *P* = 4.0×10^−11^) is an intron variant in *TCF7L2*^5^. Variant rs527320094 (OR= 2.74, *P* = 4.6×10^−9^), located in *LOC105378797*, is also novel rare-variant association, with no other significant SNP (either common or rare) within a ±1 Mb window. We did not observe any substantial difference in association signals for these four variants between the results from BOLT-LMM^37^ and logistic regression considering the difference in sample size (**Supplementary Table 6**).

### Sex or age heterogeneity analysis

To examine sex or age heterogeneity in the SNP effects, we performed a GWAS analysis within each sex (male or female) and by age (two age categories separated by the median year of birth) in UKB and tested the difference in the estimated SNP effects between the two sex (or age) groups using a heterogeneity test (**Supplementary Note 6**). There was no evidence for sex heterogeneity (**Supplementary Fig. 6**), consistent with the observation that the male-female genetic correlation estimated by bivariate LDSC^23^ was not significantly different from 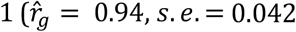, and *P*_difference_ = 0.17). There was only one genome-wide significant signal (rs72805579 at the *TMEM17* locus with *P*_heterogeneity_ = 2.1×10^−9^) with age heterogeneity (**Supplementary Fig. 6**). The estimates of SNP effects were of opposite directions in the two age groups, but the effect was not genome-wide significant in either age group (**Supplementary Table 7**). In addition, the

### Gene expression and DNA methylation associated with T2D

Most previous studies have reported the gene in closest physical proximity to the most significant SNP at a GWAS locus. However, gene regulation can be influenced by genetic variants that are physically distal to the genes^38^. To prioritize genes identified through the genome-wide significant loci that are functionally relevant to the disease, we performed an SMR analysis^39^ to test for association between the expression level of a gene and T2D using summary data of GWAS from our meta-analysis and expression quantitative trait loci (eQTL) from the eQTLGen (*n* = 14,115) and CAGE consortia (*n* = 2,765)^40^ (**Methods**). In both eQTL data sets, gene expression levels were measured in blood, and the cis-eQTL within 2 Mb of the gene expression probes with *P*_eQTL_ < 5×10^−8^ were selected as the instrumental variables in the SMR test. We identified 40 genes in eQTLGen and 24 genes in CAGE at an experimental-wise significance level (*P*_SMR_ < 2.7×10^−6^, *i.e.*, 0.05/*m*_SMR_, with *m*_SMR_ = 18,602 being the total number of SMR tests in the two data sets) (**Supplementary Tables 8-9**). To filter out the SMR associations due to linkage (*i.e.*, two causal variants in LD, one affecting gene expression and the other affecting T2D risk), all the significant SMR associations were followed by a HEIDI^39^ (HEterogeneity In Dependent Instruments) test implemented in the SMR software tool (**Methods**). Therefore, genes not rejected by HEIDI were those associated with T2D through pleiotropy at a shared genetic variant. Of the genes that passed the SMR test, 27 genes in eQTLGen and 15 genes in CAGE were not rejected by the HEIDI test (*P*_HEIDI_ > 7.8×10^−4^, *i.e.*, 0.05/*m*_SMR_, with *m*_SMR_ = 64 being the total number of SMR tests in the two data sets) (**Table 2** and **Supplementary Tables 8-9**), with seven genes in common and 33 unique genes in total. SNPs associated with the expression levels of genes including *EHHADH* (rs7431357), *SSSCA1* (rs1194076) and *P2RX4* (rs2071271) in eQTLGen were not significant in the T2D meta-analysis, likely because of the lack of power; these SNPs are expected to be detected in future studies with larger sample sizes.

To identify the regulatory elements associated with T2D risk, we performed SMR analysis using methylation quantitative trait locus (mQTL) data from McRae *et al*.^41^ (*n* = 1,980) to identify DNA methylation (DNAm) sites associated with T2D through pleiotropy at a shared genetic variant. In total, 235 DNAm sites were associated with T2D, with *P*_SMR_ < 6.3×10^−7^ (*m*_SMR_ = 78,961) and *P*_HEIDI_ > 1.6×10^−4^ (*m*_HEIDI_ = 323) (**Supplementary Table 10**); these sites were significantly enriched in promoters (fold change = 1.60 and *P*_enrichment_ = 1.6×10^−7^) and weak enhancers (fold change = 1.74 and *P*_enrichment_ = 1.4×10^−2^) (**Supplementary Note 7** and **Supplementary Fig. 7**). Identification of DNAm sites and their target genes relies on consistent association signals across omics levels^15^. To demonstrate this, we conducted the SMR analysis to test for associations between the 235 T2D-associated DNAm sites and the 33 T2D-associated genes and identified 22 DNAm sites associated with 16 genes in eQTLGen (**Supplementary Table 11**) and 21 DNAm sites associated with 15 genes in CAGE (**Supplementary Table 12**) at *P*_SMR_ < 2.5×10^−7^ (*m*_SMR_ = 202,609) and *P*_HEIDI_ > 2.1×10^−4^ (*m*_HEIDI_ = 235). These results can be used to infer plausible regulatory mechanisms for how genetic variants affect T2D risk by regulating the expression levels of genes through DNAm (see below).

### SMR associations in multiple T2D-relevant tissues

To replicate the SMR associations in a wider range of tissues relevant to T2D, we performed SMR analyses based on cis-eQTL data from four tissues in GTEx (*i.e.*, adipose subcutaneous tissue, adipose visceral omentum, liver and pancreas). We denoted these four tissues as GTEx-AALP. Of the 27 putative T2D genes identified by SMR and HEIDI using the eQTLGen data, 10 had a cis-eQTL at *P*_eQTL_ < 5×10^−8^ in at least one of the four GTEx-AALP tissues (**Supplementary Table 13**). Note that the decrease in eQTL detection power is expected given the much smaller sample size of GTEx-AALP (*n* = 153 to 385) compared to that of eQTLGen (*n* = 14,115). As a benchmark, 17 of the 27 genes had a cis-eQTL at *P*_eQTL_ < 5×10^−8^ in GTEx blood (*n* = 369). We first performed the SMR analysis in GTEx-blood and found that 12 of the 17 genes were replicated at *P*_SMR_ < 2.9×10^−3^ (*i.e.*, 0.05 / 17) (**Supplementary Table 13**). We then conducted the SMR analysis in GTEx-AALP. The result showed that 8 of the 10 genes showed significant SMR associations at *P*_SMR_ < 1.3×10^−3^ (*i.e.*, 0.05 / (10 × 4)) in at least one of the four GTEx-AALP tissues, a replication rate comparable to that found in GTEx-blood. Among the 8 genes, *CWF19L1*, for which the cis-eQTL effects are highly consistent across different tissues, was significant in all the data sets (**Supplementary Fig. 8**).

The replication analysis described above depends heavily on the sample sizes of eQTL studies. A less sample-size-dependent approach is to quantify how well the effects of the top associated cis-eQTLs for all the 27 putative T2D genes estimated in blood (*i.e.*, the eQTLGen data) correlate with those estimated in the GTEx tissues, accounting for sampling variation in estimated SNP effects^42^. This approach avoids the need to use a stringent *P*-value threshold to select cis-eQTLs in the GTEx tissues with small sample sizes. We found that the mean correlation of cis-eQTL effects between eQTLGen blood and GTEx-AALP was 0.47 (*s.e.* = 0.16), comparable to and not significantly different from the value of 0.64 (*s.e.* = 0.16) between eQTLGen and GTEx blood. We also found that the estimated SMR effects of 18 genes that passed the SMR test and were not rejected by the HEIDI test in either eQTLGen or GTEx were highly correlated (Pearson’s correlation *r* = 0.80) (**Supplementary Fig. 9**). Note that this correlation is not expected to be unity because of differences in the technology used to measure gene expression (Illumina gene expression arrays for eQTLGen vs. RNA-seq for GTEx).

These results support the validity of using eQTL data from blood for the SMR and HEIDI analysis; using this method, we can make use of eQTL data from very large samples to increase the statistical power, consistent with the conclusions of a recent study^42^. In addition, blood is also considered to be a T2D-relevant tissue, and tissue-specific effects that are not detected in blood will affect the power of the SMR and HEIDI analysis rather than generating false positive associations.

### Putative regulatory mechanisms for three T2D genes

Here, we use the genes *CAMK1D, TP53INP1* and *ATP5G1* as examples to hypothesize possible mechanisms of how genetic variants affect T2D risk by controlling DNAm for gene regulation^15^. Functional gene annotation information was acquired from the Roadmap Epigenomics Mapping Consortium^43^ (REMC).

The significant SMR association of *CAMK1D* with T2D was identified in both eQTL data sets (**Table 2** and **Supplementary Tables 10-11**). The top eQTL, rs11257655, located in the intergenic region (active enhancer) between *CDC123* and *CAMK1D*, was also a genome-wide significant SNP in our meta-analysis (*P* = 2.0×10^−17^). It was previously shown that rs11257655 is located in the binding motif for *FOXA1*/*FOXA2* and that the T allele of this SNP is a risk allele that increases the expression level of *CAMK1D* through allelic-specific binding of *FOXA1* and *FOXA2*^44^. Another functional study demonstrated that increasing the expression of *FOXA1* and its subsequent binding to enhancers was associated with DNA demethylation^45^. Our analysis was consistent with previous studies in showing that the T allele of rs11257655 increases both *CAMK1D* transcription (*β* = 0.553 and *s.e.* = 0.014, where *β* is the allele substitution effect on gene expression in standard deviation units) and T2D risk (OR = 1.076 and *s.e.* = 0.009) (**Supplementary Tables 8-9, 11**). Moreover, rs11257655 was also the top mQTL (**Fig. 2**); the T allele of this SNP is associated with decreased methylation at the site cg03575602 in the promoter region of *CAMK1D*, suggesting that the T allele of rs11257655 up-regulates the transcription of *CAMK1D* by reducing the methylation level at cg03575602. Leveraging all the information above, we proposed the following model of the genetic mechanism at *CAMK1D* for T2D risk (**Fig. 3**). In the presence of the T allele at rs11257655, *FOXA1*/*FOXA2* and other transcription factors bind to the enhancer region and form a protein complex that leads to a decrease in the DNAm level of the promoter region of *CAMK1D* and recruits the RNA polymerase to the promoter, resulting in an increase in the expression of *CAMK1D* (**Fig. 3**). A recent study showed that the T risk allele is correlated with reduced DNAm and increased chromatin accessibility across multiple islet samples^46^ and that it is associated with disrupted beta cell function^47^. Our inference highlights the role of promoter-enhancer interaction in gene regulation; this interaction was analytically indicated by the integrative analysis using the SMR and HEIDI approaches.

**Figure 2.**
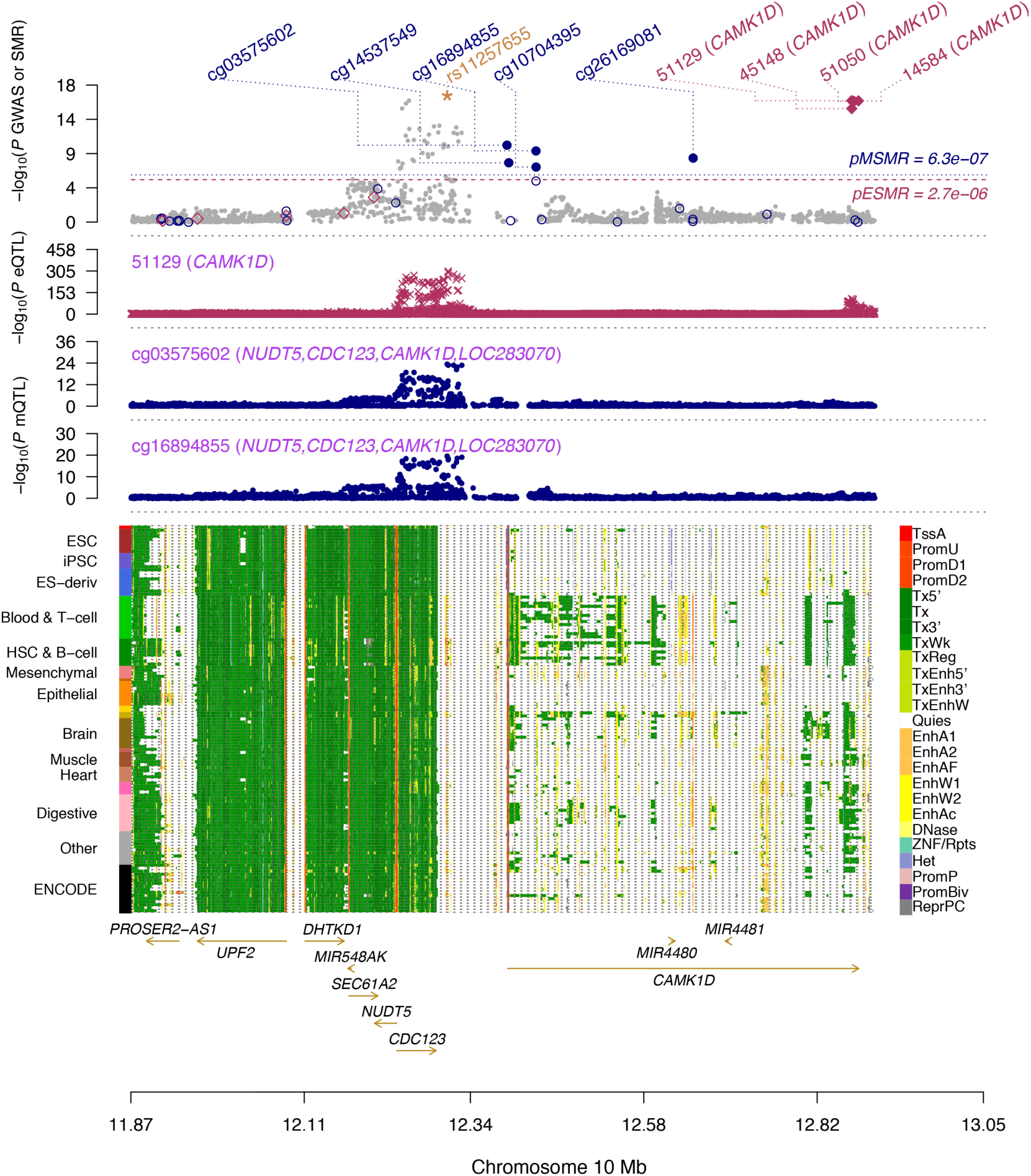
Prioritizing genes and regulatory elements at the *CDC123*/*CAMK1D* locus for T2D. The results of the SMR analysis that integrates data from GWAS, eQTL and mQTL studies are shown. The top plot shows −log10(*P*-value) of SNPs from the GWAS meta-analysis for T2D. Red diamonds and blue circles represent −log10(*P*-value) from the SMR tests for associations of gene expression and DNAm probes with T2D, respectively. Solid diamonds and circles represent the probes not rejected by the HEIDI test. The yellow star denotes the top cis-eQTL SNP rs11257655. The second plot shows −log10(*P*-value) of the SNP association for gene expression probe 51129 (tagging *CAMK1D*). The third plot shows −log10(*P*-value) of the SNP association with DNAm probes cg03575602 and cg16894855 from the mQTL study. The bottom plot shows 25 chromatin state annotations (indicated by colors) of 127 samples from Roadmap Epigenomics Mapping Consortium (REMC) for different primary cells and tissue types (rows).

**Figure 3.**
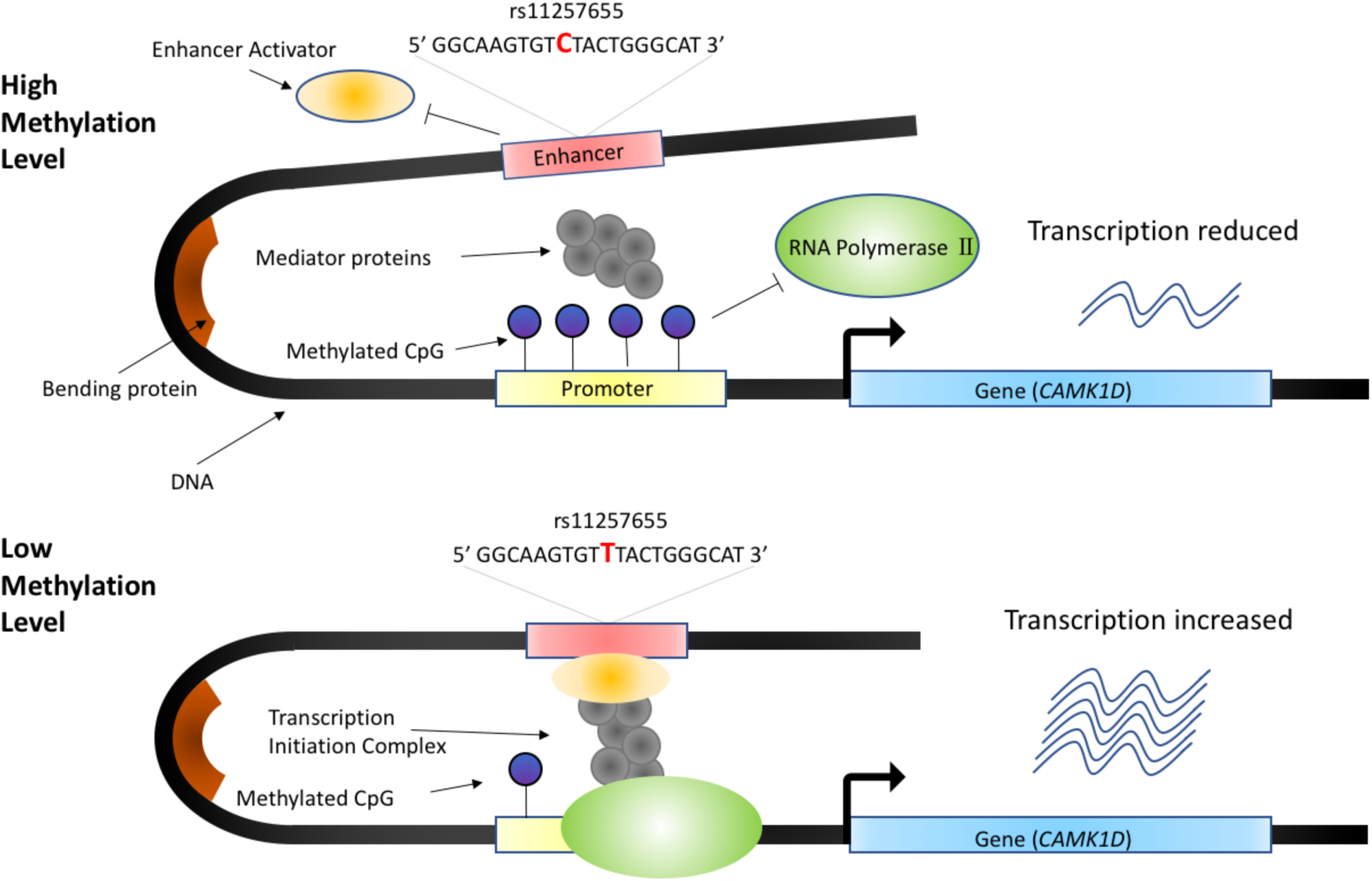
Hypothesized mechanism of how a *CAMK1D* variant affects T2D risk. When the allele of rs11257655 in the enhancer region (red) changes from C to T, the enhancer activator protein *FOXA1*/*FOXA2* (orange ellipsoid) binds to the enhancer region and the DNA methylation level in the promoter region is reduced; this increases the binding efficiency of RNA polymerase II recruited by mediator proteins (gray circles) and therefore increases the transcription of *CAMK1D*.

The second example is *TP53INP1*, the expression level of which is positively associated with T2D as indicated by the SMR analysis (**Table 2**). This is supported by previous findings that the protein encoded by *TP53INP1* regulates the *TCF7L2*-p53-p53INP1 pathway in such a way as to induce apoptosis and that the survival of pancreatic beta cells is associated with the level of expression of *TP53INP1*^48^. *TP53INP1* was mapped as the target gene for three DNAm sites (cg13393036, cg09323728 and cg23172400) by SMR (**Fig. 4**). All three DNAm sites were located in the promoter region of *TP53INP1* and had positive effects on the expression level of *TP53INP1* and on T2D risk (**Supplementary Tables 7** and **9-10**). Based on these results, we proposed the following hypothesis for the regulatory mechanism (**Fig. 5**). When the DNAm level of the promoter region is low, expression of *TP53INP1* is suppressed due to the binding of repressor(s) to the promoter. When the DNAm level of the promoter region is high, the binding of repressor(s) is disrupted, allowing the binding of transcription factors that recruit RNA polymerase and resulting in up-regulation of gene expression. Increased expression of this gene has been shown to increase T2D risk by decreasing the survival rate of pancreatic beta cells through a *TCF7L2*-p53-p53INP1-dependent pathway^49,50^.

**Figure 4.**
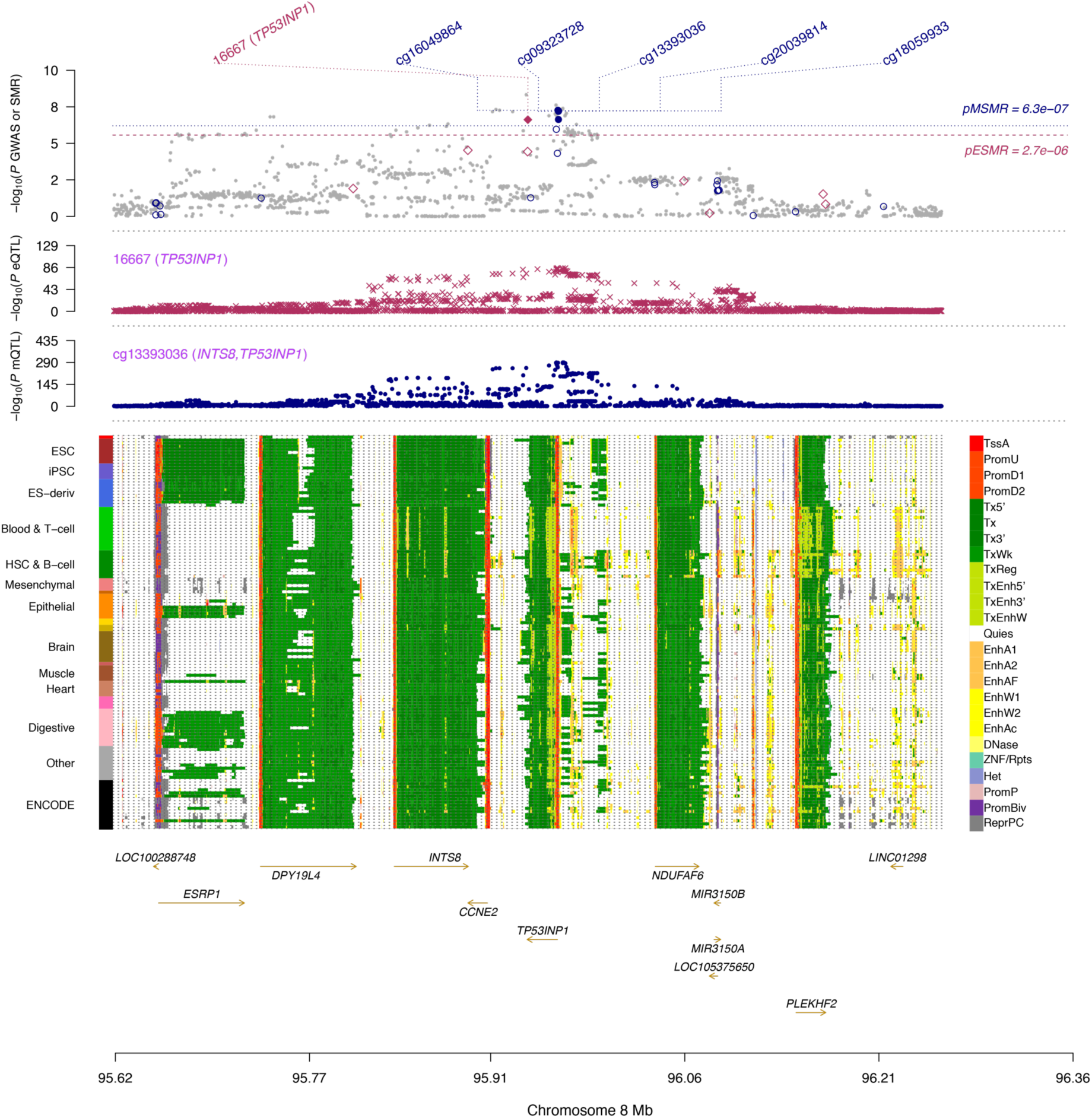
Prioritizing genes and regulatory elements at *TP53INP1* locus for T2D. Shown are the results from the SMR analysis that integrates data from GWAS, eQTL and mQTL studies. The top plot shows −log10(*P*-value) from the GWAS meta-analysis for T2D. Red diamonds and blue circles represent −log10(*P*-value) from the SMR tests for associations of gene expression and DNAm probes with T2D, respectively. Solid diamonds and circles represent the probes not rejected by the HEIDI test. The second plot shows −log10(*P*-value) of the SNP association with gene expression probe 16667 (tagging *TP53INP1*). The third plot shows −log10(*P*-value) of the SNP association with DNAm probe cg13393036 and cg09323728. The bottom plot shows 25 chromatin state annotations (indicated by colors) of 127 samples from Roadmap Epigenomics Mapping Consortium (REMC) for different primary cells and tissue types (rows).

**Figure 5.**
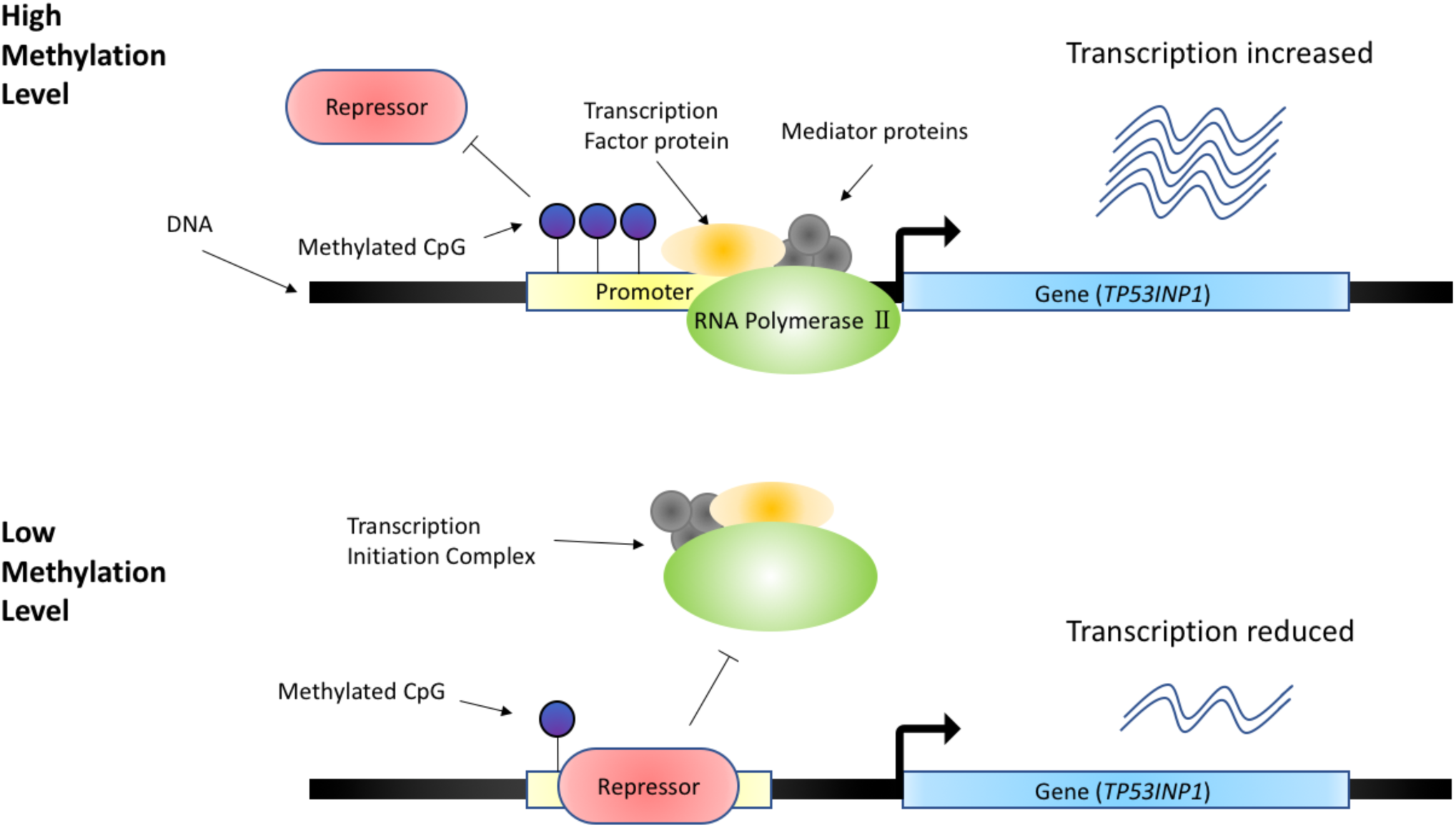
Hypothesized mechanism of how *TP53INP1* affects T2D risk. When the promoter region is highly methylated, which prevents binding of repressor protein (red rounded rectangle) to the promoter region, RNA polymerase II (green ellipsoid), transcription factor protein (orange ellipsoid) and mediator proteins (gray circles) will form a transcription initiation complex that increases the transcription. However, when the methylation level of the promoter region is low, repressor protein can more efficiently bind to the promoter, blocking the binding of the transcription initiation complex to the promoter, which decreases the transcription of *TP53INP1*.

The third example involves two proximal genes, *ATP5G1* and *UBE2Z*, the expression levels of which were significantly associated with T2D according to the SMR analysis (**Table 2**). A methylation probe (cg16584676) located in the promoter region of *UBE2Z* was associated with the expression levels of both *ATP5G1* and *UBE2Z* (**Supplementary Fig. 10a**), suggesting that these two genes are co-regulated by a genetic variant through DNAm. The effect of cg16584676 on gene expression was negative (**Supplementary Tables 10-11**), implying the following plausible mechanism. A genetic variant near *ATP5G1* exerts an effect on T2D by increasing the DNAm levels of the promoters for *ATP5G1* and *UBE2Z*; this decreases the binding efficiency of the transcription factors that recruit RNA polymerase, resulting in down-regulation of gene expression and ultimately leading to an increase in T2D risk (**Supplementary Fig. 10b**). *ATP5G1* has been shown to encode a subunit of mitochondrial ATP synthase, and *UBE2Z* is a ubiquitin-conjugating enzyme. Insulin receptors could be degraded by *SOCS* proteins during ubiquitin-proteasomal degradation, and *ATP5G1* and *UBE2Z* are likely to be involved in this pathway^51^. The function of insulin receptors is to regulate glucose homeostasis through the action of insulin and other tyrosine kinases, and dysfunction of these receptors leads to insulin resistance and increases T2D risk. Interestingly, in addition to cg16584676, there were four other DNAm sites in the vicinity that were significantly associated with T2D (passed SMR and not rejected by HEIDI). These four methylation sites are located in the promoter regions of *ATP5G1* (cg11715999), *GIP* (cg20551517) and *SNF8* (cg26022315 and cg07967210). *GIP* has been reported to be associated with T2D^52^. *SNF8* is a component of a complex that regulates ubiquitin-proteasomal degradation. These observations imply that these four genes (*ATP5G1, UBE2Z, GIP* and *SNF*) are probably co-expressed through promoter-promoter interactions.

The three examples above provide hypotheses for how genetic variants may affect T2D risk through regulatory pathways and demonstrate the power of integrative analysis of omics data for this purpose. These examples describe putative candidates that could be prioritized in future functional studies.

### Potential drug targets

In the SMR analysis described above, we identified 33 putative T2D genes. We matched these genes in the DrugBank database (**URLs**) and found that three genes (*ARG1, LTA* and *P2RX4*) are the targets of several approved drugs (drugs that have been approved in at least one jurisdiction). *ARG1* (UniProt ID: P05089), whose expression level was negatively associated with T2D risk, is targeted by three approved drugs: ornithine (DrugBank ID: DB00129), urea (DrugBank ID: DB03904) and manganese (DrugBank ID: DB06757), but the pharmacological mechanism of action of these drugs remains unknown. Arginase (*ARG1* is an isoform in liver) is a manganese-containing enzyme that catalyzes the hydrolysis of arginine to ornithine and urea. Arginase in vascular tissue might be a potential therapeutic target for the treatment of vascular dysfunction in diabetes^53^. Metformin, an oral antidiabetic drug that is used in the treatment of diabetes, was reported to increase *ARG1* expression in a murine macrophage cell line^54^, consistent with our SMR result that increased expression of *ARG1* is associated with decreased T2D risk (**Supplementary Table 8**). There is also evidence for an interaction between *ARG1* and metformin (Comparative Toxicogenomics Database, **URLs**). The likely mechanism is that metformin activates AMP-activated protein kinase (AMPK), resulting in increased expression of *ARG1*^55^, again consistent with our SMR result. *LTA* (UniProt ID: P08637), whose expression level was negatively associated with T2D risk, is targeted by the approved drug etanercept (DrugBank ID: DB00005) for rheumatoid arthritis (RA) treatment. Previous studies have shown that genetic variants in the *LTA*-*TNF* region are significantly associated with the response of early RA to etanercept treatment^56,57^. *P2RX4* (UniProt ID: Q99571), the expression level of which was positively associated with T2D risk, is targeted by eslicarbazepine acetate (DrugBank ID: DB09119; antagonist for *P2RX4*). Eslicarbazepine acetate is an anticonvulsant that inhibits repeated neuronal firing and stabilizes the inactivated state of voltage-gated sodium channels; its pharmacological action makes it useful as an adjunctive therapy for partial-onset seizures^58^. Antagonists of *P2RX4* inhibit high glucose, prevent endothelial cell dysfunction^59^, and are useful in the treatment of diabetic nephropathy^60^.

To explore whether any of these three genes have potential adverse effects, we checked the associations of the lead variants at the three loci with other traits from previous studies, including two insulin-related GWAS (insulin sensitivity^30^ and insulin secretion^29^) and four lipid traits (HDL cholesterol, LDL cholesterol, triglycerides and total cholesterol)^61^ (**Supplementary Table 14**). We did not observe any significant association with insulin traits after correcting for multiple testing (*i.e.*, 0.05 / (3 × *t*), where *t* is the number of traits). However, the risk allele of the lead T2D-associated variant at the *LTA* locus was associated with increased LDL cholesterol, total cholesterol and triglycerides. The risk allele of the lead T2D-associated variant at the *ARG1* locus was associated with decreased HDL cholesterol and total cholesterol.

In addition to the three genes that are currently targeted by approved drugs, we found two additional genes that are targeted by an approved veterinary drug and a nutraceutical drug, respectively (**URLs** and **Supplementary Note 8**).

### Enrichment of genetic variation in functional regions and tissue/cell types

Recent studies have indicated that different functional regions of the genome contribute disproportionately to total heritability^62^. We applied a stratified LD score regression method^62^ to dissect the contributions of the functional elements to the SNP-based heritability 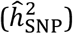 for T2D. There were significant enrichments in some functional categories (**Supplementary Fig. 11** and **Supplementary Table 15**). First, the conserved regions in mammals^63^ showed the largest enrichment, with 2.6% of SNPs explaining 24.8% of 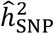 (fold-change = 9.5; *P* = 1.9×10^−4^). This supports the biological importance of conserved regions, although the functions of many conserved regions are still undefined. Second, the histone marker H3K9ac^64^ was highly enriched, with 12.6% of SNPs explaining 59.7% of 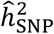 (fold-change = 4.7; *P* = 2.5×10^−5^). H3K9ac can activate genes by acetylation and is highly associated with active promoters. We also partitioned 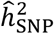 into ten cell type groups (**Supplementary Table 16**); the top cell type group for T2D was “adrenal or pancreas” (fold-change = 6.0; *P* = 8.1×10^−9^), and the result was highly significant (*P*_Bonferroni_ = 1.8×10^−6^) after Bonferroni correction for 220 tests (**Supplementary Fig. 12**).

We further used MAGMA^65^ to test the enriched gene sets. In total, 305 gene sets in GO_BP terms and 20 gene sets in KEGG pathways were significantly enriched (**Supplementary Table 17**). The top pathway enrichment was “glucose homeostasis” (*P* = 6.0×10^−8^) in GO_BP, and “maturity onset diabetes of the young” (*P* = 3.2×10^−7^) in KEGG. To further investigate the molecular connections of T2D-associated genes, a protein-protein interaction network was analyzed using STRING^66^ (**Supplementary Fig. 13**). Among the functional enrichment (**Supplementary Table 18**) in this network, there are four genes (*HHEX, HNF1A, HNF1B*, and *FOXA2*) involved in the KEGG pathway of “maturity onset diabetes of the young”, and four genes (*ADCY5, CAMK2G, KCNJ11*, and *KCNU1*) were enriched in “insulin secretion”.

### Natural selection of T2D-associated variants

We performed an LD- and MAF- stratified GREML analysis^67^ (**Methods**) in a subset of unrelated individuals in UKB (*n* = 15,767 cases and 104,233 controls) to estimate the variance explained by SNPs in different MAF ranges (*m* = 18,138,214 in total). We partitioned the SNPs into 7 MAF bins with high and low LD bins within each MAF bin to avoid MAF- and/or LD-mediated bias in 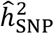 (**Methods**). The 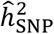 was 33.2% (*s.e.* = 2.1%) on the liability scale (**Supplementary Table 19**). Under an evolutionary neutral model and a constant population size^68^, the explained variance is uniformly distributed as a function of MAF, which means that the variance explained by variants with MAF ≤ 0.1 equals that explained by variants with MAF > 0.4. However, in our results, the MAF bin containing low-MAF and rare variants (MAF < 0.1) showed a larger estimate than any other MAF bin (**Fig. 6a** and **Supplementary Table 19**), consistent with a model of negative (purifying) selection or population expansion^69^. To further distinguish between the two models (negative selection vs. population expansion), we performed an additional analysis using a recently developed method, BayesS^70^, to estimate the relationship between variance in effect size and MAF (**Methods**). The method also allowed us to estimate 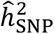 and polygenicity (*π*) on each chromosome. The results (**Fig. 6b**) showed that the 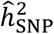 of each chromosome was highly correlated with its length (*r* = 0.92), consistent with the results of previous studies for height and schizophrenia^71,72^. The mean estimate of *π, i.e.*, the proportion of SNPs with non-zero effects, was 1.75% across all chromosomes (**Fig. 6c** and **Supplementary Table 20**), suggesting a high degree of polygenicity for T2D. The sum of per-chromosome 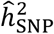 from BayesS was 31.9% (*s.e.* = 4.1%) on the liability scale, slightly higher than that based on HapMap3 SNPs from an HE regression analysis (28.7%, *s.e.* = 1.1%) using a full set of unrelated UKB individuals (*n* = 348,580) or from an LD score regression analysis (22.6%, s.e = 1.2%) using all the UKB individuals (*n* = 455,607) (**Supplementary Table 21**). The variance in effect size was significantly negatively correlated with MAF (*Ŝ*= −0.53, *s.e.* = 0.09), consistent with a model of negative selection on deleterious rare alleles (**Fig. 6d**) and inconsistent with a recent study^12^ concluding that T2D-associated loci have not been under natural selection. Our conclusion regarding negative selection is also consistent with the observation that the minor alleles of 9 of the 11 rare variants at *P* < 5E-8 were T2D risk alleles (**Supplementary Table 6**). The signal of negative selection implies that a large number of rare variants will be discovered in future GWAS in which appropriate genotyping strategies are used.

**Figure 6.**
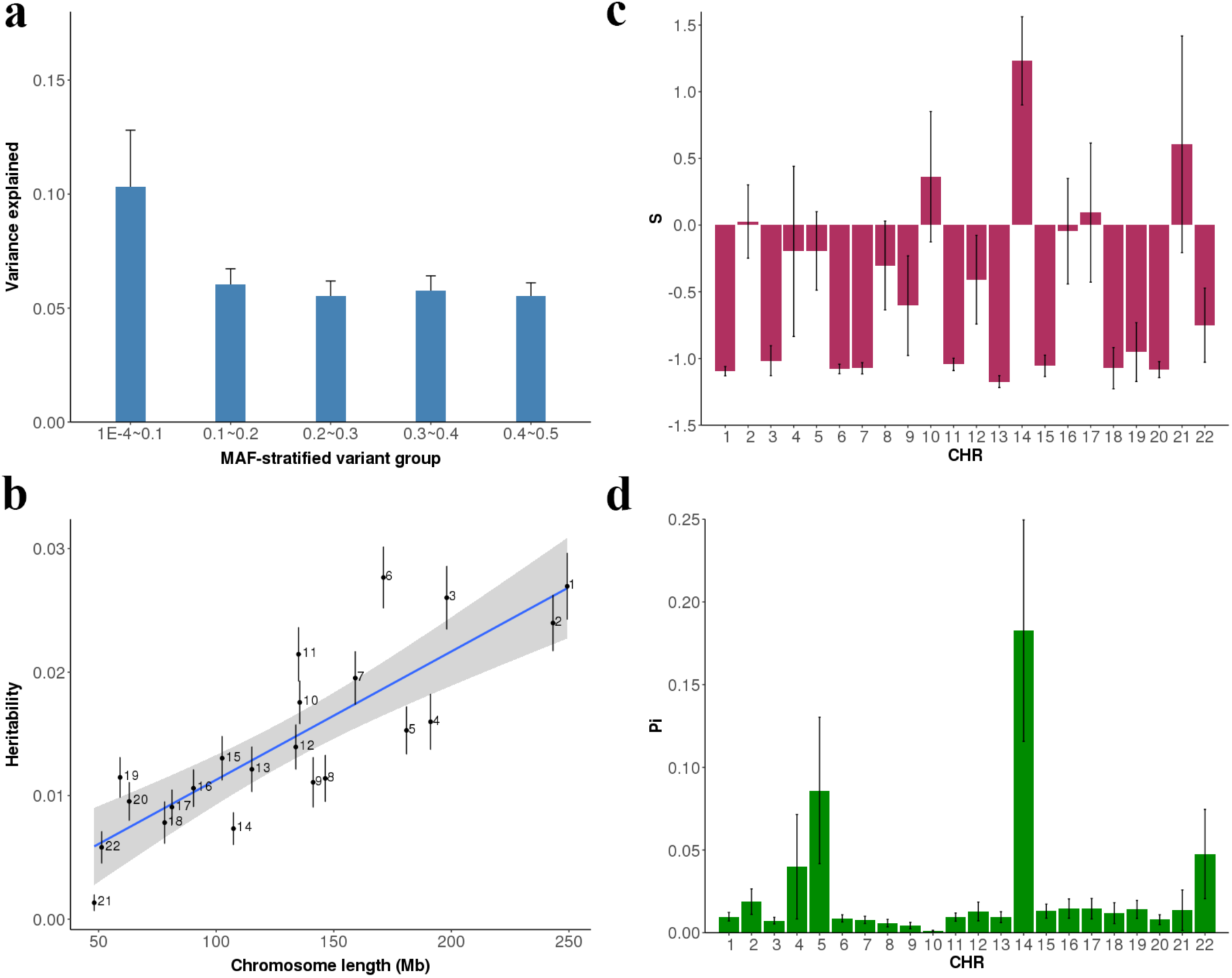
Estimating the SNP-based heritability and polygenicity and detecting signals of purifying selection in the UKB data. Shown in panel are the results from the GREML-LDMS analysis. Shown in panels b, c and d are the results from the BayesS analysis. Error bars are standard errors of the estimates. a) Variance explained by SNPs in each MAF bin. We combined the estimates of the first three bins (MAF < 0.1) to harmonize the width of all MAF bins. b) Chromosome-wide SNP-based heritability against chromosome length. c) Estimate of the BayesS parameter (*S*) reflecting the strength of purifying selection on each chromosome. d) Proportion of SNPs with non-zero effects on each chromosome (*π*).

### Polygenic risk score (PRS) analysis

We used DIAGRAM and UKB as the discovery set and GERA as a validation set in the PRS analysis^73^. To avoid sample overlap between the discovery and validation sets, we re-ran the meta-analysis excluding the GERA cohort and identified 109 near-independent common SNPs at *P* < 5×10^−8^. These SNPs were then used to derive prediction equations for individuals in GERA (**Methods**). We divided GERA into ten subsets to acquire the sampling variance of the estimated classification accuracy. On average, the classification accuracy (measured by the area under the curve or AUC^74^) was 0.579 (*s.e.* = 0.003), lower than the classification accuracy of 0.599 (*s.e.* = 0.002) obtained using all SNP effects (∼5.1 million SNPs) estimated from GCTA-SBLUP (Summary-data-based Best Linear Unbiased Prediction)^75^ (**Supplementary Table 22**). We further quantified the variance explained by the 109 genome-wide significant SNPs by fitting them to a multiple regression model with phenotypes in GERA. These SNPs explained 3.9% of the phenotypic variance on the liability scale compared with an estimate of 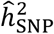 of 7.2% from GREML using HapMap3 SNPs, although the 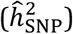 in GERA was much lower than that in UKB.

## Discussion

In this study, we sought to identify novel genetic loci associated with T2D by a meta-analysis of GWAS with the largest sample size to date and to infer plausible genetic regulation mechanisms at known and novel loci by an integrative analysis of GWAS and omics data. We identified 139 near-independent common variants (*P* < 5 × 10^−8^) and 4 rare variants (*P* < 5 × 10^−9^) for T2D in the meta-analysis. Of the 139 common loci, 39 were novel compared with the results of all 49 previous T2D GWAS from the GWAS Catalog (**URLs**)^76^, including the two recent studies by DIAGRAM^52^ and Zhao *et al*.^77^. By integrating omics data, we have inferred the genetic mechanisms for the three genes *CAMK1D, TP53INP1* and *ATP5G1*; the inferred mechanisms suggest that enhancer-promoter interactions with DNA methylation play an important role in mediating the effects of genetic variants on T2D risk. These findings provide deeper insight into the etiology of T2D and suggest candidate genes for functional studies in the future. Furthermore, our estimation of genetic architecture suggests that T2D is a polygenic trait for which both rare and common variants contribute to the genetic variation and indicates that rarer variants tend to have larger effects on T2D risk (**Fig. 7**). Assuming that most new mutations are deleterious for fitness, our result is consistent with a model in which mutations that have larger effects on T2D (and thereby on fitness through pleiotropy) are more likely to be maintained at low frequencies in the population by purifying selection.

This study has a number of limitations. First, the SNP-T2D associations identified by the meta-analysis might be biased by misdiagnosis of T1D (type 1 diabetes) and LADA (latent autoimmune diabetes in adults)^78^. Previous studies found that biases in SNP-T2D associations due to misdiagnosis were likely to be very modest^7,52,79^. We showed by two additional analyses based on known T1D loci that most of the novel SNP-T2D associations identified in this study are unlikely to be driven by misdiagnosed T1D cases (**Supplementary Note 9** and **Supplementary Table 23**). Second, some of the T2D-associated SNPs might confer T2D risk through mediators such as obesity or dyslipidemia. To explore this possibility, we performed a summary-data-based conditional analysis of the 139 T2D-associated SNPs conditioning on body mass index (BMI) or dyslipidemia by GCTA-mtCOJO^80^ using GWAS data for these two traits from UKB. It appeared that the effect sizes of most T2D-associated SNPs, with the exception of a few outliers (e.g., *FTO, MC4R, POCS* and *TFAP2B*), were not affected by BMI or dyslipidemia (**Supplementary Fig. 14**). These loci were among those showing the strongest associations with BMI^81^, consistent with the finding from a previous T2D study^82^. Third, among the 39 novel loci, there was only one locus (*ARG1*/*MED23*, **Supplementary Fig. 15**) at which the association between gene expression and T2D risk was significant in SMR and not rejected by HEIDI (**Table 2**). This is because the power of the SMR test depends primarily on the SNP effect from GWAS^13^, which is small for the novel loci. Finally, we employed the SMR and HEIDI methods to map CpG sites to their target genes and to identify the CpG sites associated with T2D because of pleiotropy. The SMR approach uses genome-wide significant mQTL as an instrumental variable for each CpG site, which requires a large sample size for the mQTL discovery. In this study, we used mQTL data based on Illumina HumanMethylation450 arrays because of the relatively large sample size (*n* = 1,980). Unfortunately, we did not have access to mQTL data from whole-genome bisulfite sequencing (WGBS) in a large sample. Nevertheless, it is noteworthy that there are three T2D-associated variants at the *CAMK1D*/*CDC123, ADCY5*, and *KLHDC5* loci that show hypomethylation and allelic imbalance as identified by Thurner *et al*.^46^ using WGBS data (*n* = 10), all of which were genome-wide significant in our mQTL-based SMR analysis. Despite these limitations, our study highlights the benefits of integrating multiple omics data to identify functional genes and putative regulatory mechanisms driven by local genetic variation. Future applications of integrative omics data analyses are expected to increase our understanding of the biological mechanisms underlying human disease.

## Methods

### Summary statistics of DIAGRAM, GERA, and UKB

The data used in this study were derived from 659,316 individuals of European ancestry and a small cohort from Pakistan and were obtained from three data sets: DIAbetes Genetics Replication And Meta-analysis (DIAGRAM)^7^, Genetic Epidemiology Research on Adult Health and Aging (GERA)^16^ and UK Biobank (UKB)^17^.

DIAGRAM: The DIAGRAM data were obtained from publicly available databases (**URLs**) and included two stages of summary statistics. In stage 1, there were 12,171 cases and 56,862 controls from 12 GWAS cohorts of European descent, and the genotype data were imputed to the HapMap2 Project^83^ (∼2.5 million SNPs after quality control). In stage 2, there were 22,669 cases and 58,119 controls genotyped on Metabochips (∼137,900 SNPs), including 1,178 cases and 2,472 controls of Pakistani descent. There was limited evidence of genetic heterogeneity between individuals of European and those of Pakistani descent for T2D^7^. The sample prevalence was 23.3% (17.6% in stage 1 and 28.1% in stage 2). We imputed the stage 1 summary statistics by ImpG^19^ and combined the imputed data with stage 2 summary statistics (**Supplementary Note 1**).

GERA: There were 6,905 cases and 46,983 controls in GERA, and the sample prevalence was 12.4%. We cleaned the GERA genotype data using standard quality control (QC) filters (excluding SNPs with missing rate ≥ 0.02, Hardy-Weinberg equilibrium test *P*-value ≤ 1×10^−6^ or minor allele count ≤ 1 and removing individuals with missing rate ≥ 0.02) and imputed the genotype data to the 1000 Genomes Projects (1KGP) reference panels^84^ using IMPUTE2^85^. We used GCTA^86^ to compute the genetic relationship matrix (GRM) of all the individuals based on a subset of imputed SNPs (HapMap3 SNPs with MAF ≥ 0.01 and imputation info score ≥ 0.3), removed the related individuals at a genetic relatedness threshold of 0.05, and retained 53,888 individuals (6,905 cases and 46,983 controls) for further analysis. We computed the first 20 principal components (PCs) from the GRM. The summary statistics in GERA were obtained from a GWAS analysis using PLINK2^87^ with sex, age, and the first 20 principal components (PCs) fitted as covariates. To examine the influence of imputation panel on the meta-analysis result, we further imputed GERA to the Haplotype Reference Consortium^21^ (HRC) using the Sanger imputation service (**URLs**).

UKB: Genotype data from UKB were cleaned and imputed to HRC by the UKB team^17,21^. There were 21,147 cases and 434,460 controls, and the sample prevalence was 5.5%. We identified a European subset of UKB participants (*n* = 456,426) by projecting the UKB participants onto the 1KGP PCs. Genotype probabilities were converted to hard-call genotypes using PLINK2^87^ (--hard-call 0.1), and we excluded SNPs with minor allele count < 5, Hardy-Weinberg equilibrium test *P*- value < 1×10^−6^, missing genotype rate > 0.05, or imputation info score < 0.3. The UKB phenotype was acquired from self-report, ICD10 main diagnoses and ICD10 secondary diagnoses (field IDs: 20002, 41202 and 41204). The GWAS analysis in UKB was conducted in BOLT-LMM^37^ with sex and age fitted as covariates. In the BOLT-LMM analysis, we used 711,933 SNPs acquired by LD pruning (*r*^2^ < 0.9) from Hapmap3 SNPs to control for relatedness, population stratification and polygenic effects. We transformed the effect size from BOLT-LMM on the observed 0-1 scale to the odds ratio (OR) using LMOR^88^.

### Inverse variance based meta-analysis

Before conducting the meta-analysis, we performed several analyses in which we examined genetic heterogeneity and sample overlap among data sets (**Supplementary Note 2**). We performed a two-stage meta-analysis. The first stage combined DIAGRAM stage 1 (GWAS chip) data with GERA and UKB. The second stage combined DIAGRAM stage 1 and 2 (GWAS chip and metabolism chip) with GERA and UKB. We extracted the SNPs common to the three data sets (5,526,193 SNPs in stage 1 and 5,053,015 million SNPs in stage 2) and performed the meta-analyses using an inverse-variance based method in METAL^20^. The stage 1 meta-analysis data were only used to estimate the SNP-based heritability, and the stage 2 meta-analysis data were used in the follow-up analyses.

### Summary-data-based Mendelian Randomization (SMR) analysis

We performed an SMR and HEIDI analysis^39^ to identify genes whose expression levels were associated with a trait due to pleiotropy using summary statistics from GWAS and eQTL/mQTL studies. The HEIDI test^39^ uses multiple SNPs in a cis-eQTL region to distinguish pleiotropy from linkage. In the SMR analysis, we used eQTL summary data from the eQTLGen Consortium (*n* = 14,115 in whole blood), the CAGE (*n* = 2,765 in peripheral blood)^40^ and the GTEx v7 release (*n* = 385 in adipose subcutaneous tissue, *n* = 313 in adipose visceral omentum, *n* = 153 in liver, *n* = 220 in pancreas and *n* = 369 from whole blood)^89^. In CAGE and eQTLGen, gene expression levels were measured using Illumina gene expression arrays; in GTEx, gene expression levels were measured by RNA-seq. The SNP genotypes in all three cohorts were imputed to 1KGP. The mQTL summary data were obtained from genetic analyses of DNA methylation measured on Illumina HumanMethylation450 arrays (*n* = 1,980 in peripheral blood)^41^.

### Estimating the genetic architecture for T2D

The MAF- and LD-stratified GREML (GREML-LDMS) is a method for estimating SNP-based heritability that is robust to model misspecification^67,90^. For ease of computation, we limited the analysis to a subset of unrelated UKB individuals (15,767 cases and 104,233 controls); in this subset, we kept all 15,767 cases among the unrelated individuals to maximize the sample size of cases and randomly selected 104,233 individuals from 332,813 unrelated controls. We first estimated the segment-based LD score, stratified ∼18 million SNPs into two groups based on the segment-based LD scores (high vs. low LD groups), and then stratified the SNPs in each LD group into seven MAF bins (1E-4-1E-3, 1E-3-1E-2, 1E-2-0.1, 0.1-0.2, 0.2-0.3, 0.3-0.4 and 0.4-0.5). We computed the GRMs using the stratified SNPs and performed GREML analysis fitting 14 GRMs (with sex, age, and the first 10 PCs fitted as covariates) in one model to estimate the SNP-based heritability in each MAF bin. We used 10% as the population prevalence to convert the estimate to that on the liability scale.

We used GCTB-BayesS^70^ to estimate the joint distribution of SNP effect size and allele frequency. This analysis is based on 348,580 unrelated individuals (15,767 cases and 332,813 controls) and HapMap3 SNPs (∼1.23 million) with sex, age and the first 10 PCs fitted as covariates. Each SNP effect has a mixture prior of a normal distribution and a point mass at zero, with an unknown mixing probability, *π*, representing the degree of polygenicity. The variance in effect size is modeled to be dependent on MAF through a parameter S. Under an evolutionarily neutral model, SNP effect sizes are independent of MAF, *i.e., S* = 0. A negative (positive) value of *S* indicates that variants with lower MAF are prone to having larger (smaller) effects, consistent with a model of negative (positive) selection. A Markov-chain Monte Carlo (MCMC) algorithm was used to draw posterior samples for statistical inference. The posterior mean was used as the point estimate, and the posterior standard error was approximated by the standard deviation of the MCMC samples. We conducted the analysis chromosome-wise for ease of computation.

### Polygenic risk score (PRS) analysis in GERA

We used DIAGRAM and UKB as the discovery set and GERA as a validation set for the PRS analysis. To avoid sample overlap, we re-ran the meta-analysis excluding GERA and clumped significant SNPs from the meta-analysis (excluding GERA) using UKB as the reference for LD estimation (*P*- value threshold = 5×10^−8^, LD *r*^2^ threshold = 0.01 and window size = 1 Mb). After clumping, there were 109 independent SNPs. These SNPs were used to generate PRS for each individual in GERA. We then calculated the area under the curve^74^ (AUC) as a measure of classification accuracy. To quantify the sampling variance in classification accuracy, the GERA data set was divided evenly into ten groups, each with sample size ∼6,000 and similar sample prevalence. We also applied the GCTA-SBLUP (Summary-based Best Linear Unbiased Prediction) method^75^ to estimate the SNP effects when they were fitted jointly and compared the classification accuracy based on all SNPs with that based on the 109 significant SNPs.

### URLs

MAGIC consortium: https://www.magicinvestigators.org/

DrugBank: https://www.drugbank.ca/

DrugBank documentation: https://www.drugbank.ca/documentation

GWAS catalog: http://www.ebi.ac.uk/gwas/

DIAGRAM summary data: http://www.diagram-consortium.org/

Sanger imputation service: https://imputation.sanger.ac.uk/

## Supplementary Information

The supplementary information includes 10 supplementary notes, 15 supplementary figures and 23 supplementary tables.

## Contributions

J.Y., J.Z. and A.X. conceived and designed the experiment. A.X. and Y.W. performed the analysis with assistance and guidance from Z.H.Z., F.Z., L.R.L., J.S., J.Z. and J.Y. K.E.K., L.Y., ZL.Z., J.Y. and P.M.V. contributed to the analysis of the UKB data. The eQTLGen consortium provided the eQTLGen eQTL summary data. A.F.M. contributed to the analysis of DNA methylation data. A.X., J.Z. and J.Y. wrote the manuscript with the participation of all authors.

## Declaration of Interests

We declare that all authors have no competing interests.

## Data availability

Summary statistics from the meta-analysis will be available at http://cnsgenomics.com/data.html when the paper has been formally accepted for publication.

## Acknowledgments

This research was supported by the Australian National Health and Medical Research Council (1107258, 1083656, 1078037 and 1113400), Australian Research Council grants (DP160101056, DP160103860 and DP160102400), the US National Institutes of Health (R01 MH100141, P01 GM099568, R01 GM075091, R01 AG042568 and R21 ES025052), and the Sylvia & Charles Viertel Charitable Foundation. Yeda Wu is supported by the F.G. Meade Scholarship of the University of Queensland. This study makes use of data from dbGaP (accession: phs000674.v2.p2) and UK Biobank (project ID: 12505). A full list of acknowledgments of these data sets can be found in Supplementary Note 10. The members of the eQTLGen Consortium are (in alphabetical order): Mawussé Agbessi, Habibul Ahsan, Isabel Alves, Anand Andiappan, Philip Awadalla, Alexis Battle, Frank Beutner, Marc Jan Bonder, Dorret Boomsma, Mark Christiansen, Annique Claringbould, Patrick Deelen, Tõnu Esko, Marie-Julie Favé, Lude Franke, Timothy Frayling, Sina Gharib, Gregory Gibson, Gibran Hemani, Rick Jansen, Mika Kähönen, Anette Kalnapenkis, Silva Kasela, Johannes Kettunen, Yungil Kim, Holger Kirsten, Peter Kovacs, Knut Krohn, Jaanika Kronberg-Guzman, Viktorija Kukushkina, Zoltan Kutalik, Bernett Lee, Terho Lehtimäki, Markus Loeffler, Urko M. Marigorta, Andres Metspalu, Lili Milani, Martina Müller-Nurasyid, Matthias Nauck, Michel Nivard, Brenda Penninx, Markus Perola, Natalia Pervjakova, Brandon Pierce, Joseph Powell, Holger Prokisch, Bruce Psaty, Olli Raitakari, Susan Ring, Samuli Ripatti, Olaf Rotzschke, Sina Ruëger, Ashis Saha, Markus Scholz, Katharina Schramm, Ilkka Seppälä, Michael Stumvoll, Patrick Sullivan, Alexander Teumer, Joachim Thiery, Lin Tong, Anke Tönjes, Jenny van Dongen, Joyce van Meurs, Joost Verlouw, Peter Visscher, Uwe Völker, Urmo Võsa, Hanieh Yaghootkar, Jian Yang, Biao Zeng, and Futao Zhang.

