## Supplementary Materials for "Novel susceptibility loci and genetic regulation mechanisms for type 2 diabetes"

[*] Joint last author

**Contents**

**Supplementary Notes 1 to 10**

**Supplementary Figures 1 to 15**

**References**

### Supplementary Notes

#### Supplementary Note 1 Imputing the stage 1 data of DIAGRAM to 1KGP using ImpG

Since individual-level genotypes are not available in DIAGRAM, we imputed the stage 1 summary statistics of DIAGRAM to 1KGP using ImpG[1]. Before imputation, we removed SNPs on less than 9,000 cases or 50,000 controls. The haplotype reference panel files (EUR) and SNP mapping files were obtained from 1KGP phase 1 (release v3). We chose phase 1 because the ImpG-sum software does not take INDELs (insertions and deletions) into account. After removing SNPs with MAF < 0.01 or imputation accuracy metric[1] $r^2_{pred}$, 6,233,351 SNPs were retained for further analysis.

#### Supplementary Note 2 Heterogeneity and sample overlap among data sets

Before meta-analysis, we applied the LD score regression approach[2,3] to estimate the genetic correlation and sample overlap between pairwise data sets, and to assess the inflation in test-statistics in each data set. The estimates of genetic correlation between pairwise data sets were all not significantly different from 1 (**Supplementary Table 1**), suggesting the lack of evidence for genetic heterogeneity. The estimates of bivariate LD score regression intercept between pairwise data sets were all close to zero, suggesting the lack of evidence for sample overlap. We also used a metric called $\lambda_{meta}$[4] to test for sample overlap between pairwise data sets ($\lambda_{meta} > 1$ if there is sample overlap between two data sets). $\lambda_{meta}$ was 1.0 between DIAGRAM and GERA, 1.0 between DIAGRAM and UKB, and 0.99 between GERA and UKB, again suggesting the lack of evidence for sample overlap among the three data sets.

The estimate of intercept of the univariate LD score regression was 1.009 (*s.e.* = 0.008) in DIAGRAM, 1.031 (*s.e.* = 0.009) in GERA, and 1.057 (*s.e.* = 0.013) in UKB, suggesting that population stratification has been well controlled although the small inflation in GREA and UKB is worth further investigation. We performed a BOLT-LMM[5] analysis in GERA and re-ran the univariate LD score regression using the GWAS summary statistics from BOLT-LMM. The intercept was 1.026 (*s.e.* = 0.008), similar to the estimate above and still significantly larger than 1, suggesting that the small inflation in the LD score regression intercept in GERA is unlikely to be due to population structure. Note that a small inflation in LDSC intercept is often observed in data with large sample sizes[6]. Nevertheless, the remaining inflation in LDSC intercept from the BOLT-LMM summary data might be due to inflated test-statistics from a mixed linear model analysis of unbalanced case-control ratio[7].

#### Supplementary Note 3 Functional relevance of the novel gene loci to T2D

The functional relevance of some novel gene loci to T2D are supported by existing biological or molecular evidence related to insulin and glucose. For example, *MBNL1* was up-regulated by insulin stimulation[8] and controls insulin receptor (*INSR*) exon inclusion by binding to a downstream enhancer[9]. A missense mutation in *STAT3* was reported to result in neonatal diabetes by reducing insulin synthesis[10] or premature induction of pancreatic differentiation[11]. A neighbouring gene of *STAT3* (~14kb distance), *PTRF*, was reported to be associated with glucose tolerance status[12] and mediate insulin-regulated gene expression[13]. Activation of *CAMK2G* suppressed hepatic insulin signalling (insulin resistance)[14] and *FOXA2* could improve the hepatic insulin resistance in diabetic/insulin-resistant mice models[15].

#### Supplementary Note 4 GCTA-fastBAT analysis

Because the effect size of an individual SNP is often very small, it would be more powerful to detect the aggregated effect of a set of SNPs at the locus that harbours multiple associated variants. fastBAT[16] is a set-based association test approach using summary data from GWAS to test the aggregated effect of a set of SNPs within a gene[16]. We applied fastBAT to run a gene-based test using the summary-level data from the meta-analysis with LD between SNPs estimated from the 1KGP-imputed GERA data. We clustered the SNPs into 24,765 genes by physical distance and tested each gene for association at a genome-wide significance level ($P < 0.05/24{,}765 = 2.02 \times 10^{-6}$). Here, we define a novel gene discovery as a gene that passed genome-wide significance level ($P_{\text{fastBAT}} < 2.02 \times 10^{-6}$) in the gene-based analysis but there is no genome-wide significant SNP ($P_{\text{GWAS}} > 5 \times 10^{-8}$) within ±0.5 Mb of the gene.

We identified 374 genes (12 novel genes in addition to the single-SNP based meta-analysis) (**Supplementary Table 3**) for T2D at $P < 2.0 \times 10^{-6}$. The gain of power in the gene-based test can be due to the reduced multiple-testing burden or multiple independently associated variants at a locus. We therefore performed a conditional analysis at each of the 12 loci, and found that there were multiple independent signals with $P < 5 \times 10^{-5}$ at 4 of these loci (*NDUFS3*, *HIVEP2*, *ITGA1*, *FAM110D*) (**Supplementary Fig. 5**).

#### Supplementary Note 5 GCTA-COJO analysis

Conditional and joint analysis[17] (COJO) aims to identify multiple signals in a locus, conditioning on the primary associated SNP. In COJO analysis, we performed a stepwise model selection procedure to select near-independent SNPs. We set the threshold *P*-value to $5 \times 10^{-8}$, and window size of 10Mb, assuming that SNPs more than 10Mb away from each other or on different chromosomes are in linkage equilibrium. We used 1KGP-imputed GERA dataset as the reference for LD estimation. We identified 139 SNPs at the genome-wide significance threshold

(**Supplementary Table 4**). There were seven loci with multiple independent signals associated with T2D (**Supplementary Table 4**). The joint effects of the SNPs at the seven loci estimated from GCTA-COJO using summary-level data were consistent with those from multiple regression analysis of individual-level data from UKB (**Supplementary Table 5**).

**Supplementary Note 6 Sex or age heterogeneity**

We performed a GWAS analysis within each sex (male or female) or age (two age categories separated at median year of birth) group in the UKB data. In the sex heterogeneity analysis, there were 208,419 males and 247,188 females. In age heterogeneity analysis, there were 218,261 individuals in the first age group (born from 1951 to 1971) and 237,346 individuals in the second age group (born from 1930 to 1950). We then tested the difference in the estimated SNP effects between the two sex (or age) groups by a heterogeneity test, i.e. $T_d = \left(\hat{b}_1 - \hat{b}_2\right)^2/(SE_1^2 + SE_2^2)$ which follows a $\chi^2$ distribution with $df = 1$ under the null hypothesis of no difference.

**Supplementary Note 7 Enrichment of the T2D-associated DNA methylation sites in functional categories**

We obtained chromatin status data of 127 epigenomes from the Roadmap Epigenomics Mapping Consortium[18]. We mapped 235 T2D-associated DNAm sites to the 14 functional categories defined in Wu *et al.*[19] and counted the number of DNAm sites mapped to each category. We then randomly sampled from all the DNAm probes the same number of null probes with variance in DNAm levels at each probe matched and repeated the sampling 500 times. The fold enrichment value was calculated as a ratio of the observed value to that of a null probe set averaged across 500 replicates. The standard error of estimate of fold enrichment was calculated from 500 replicates (**Supplementary Fig. 7**).

**Supplementary Note 8 Additional drug targets for SMR hits**

In addition to the three genes that are currently targeted by approved drugs, we found two additional genes that are targeted by an approved veterinary drug and a nutraceutical drug (See **URLs** for detailed definition), respectively. *PLEKHA1* (UniProt ID: Q9HB21), whose expression level was negatively associated with T2D risk, is targeted by citric acid (DrugBank ID: DB04272). Intraperitoneal injection of citrate in diabetic mice reduced apoptotic and inflammatory responses and protected cardiac abnormalities induced by diabetes[20]. A reduction of citric acid cycle (CAC) flux which reflects mitochondrial dysfunction was observed in T2D patients[21]. *EHHADH* (UniProt ID: Q08426), whose expression level was negatively associated with T2D risk, is targeted by a nutraceutical drug NADH (DrugBank ID: DB00157).

**Supplementary Note 9 Biases in SNP-T2D associations due to misdiagnosed T1D or LADA cases**

Most data used in this study were from DIAGRAM and UKB. The summary statistics of DIAGRAM were from a meta-analysis of 12 GWAS cohorts. Previous studies show that the biases in SNP-T2D associations due to misdiagnosis are likely to be very modest[22-24]. Those studies claimed that their GWAS results for T2D were not confounded by T1D associations because of the absence of most known T1D-associated loci in their T2D discovery. Following the approach of Mahajan *et al.*[23], we extracted 48 T1D-associated SNPs from Bradfield *et al.*[25], and found that only four (16q23.1, *GLIS3*, 6q22.32 and *PTPN22*) of them showed associations with T2D at a suggestive significance level (i.e. $P$ < 1E-5) with two of them (*GLIS3* and 6q22.32) passing the genome-wide significance level (i.e. $P$ < 5E-8) (**Supplementary Table 23**).

Furthermore, following Scott *et al.*[24], we computed the polygenic risk score (PRS) for T1D in UKB using the 48 T1D-associated SNPs and tested for its association with the T2D phenotype. The PRS computed from the 44 T1D-only SNPs did not show significant association with T2D (OR=1.02 and $P$=0.93), unless the four risk loci that showed suggestive associations with T2D discovery were included in the computation of PRS (OR =1.99 and $P$ = 5.86E-13). These results suggest that our samples were unlikely to include a substantial number of misdiagnosed T1D cases, and that the associations of the four T1D loci with T2D were likely because of pleiotropy. We further performed the T2D GWAS by logistic regression using individual-level data with and without fitting the T1D phenotype as covariate in unrelated UKB individuals ($n$ = 347,997). The correlation coefficient of z-statistics of the 139 significant loci between the conditional and unconditional models was almost one ($r$ = 0.995). In the unconditional model, 43 out of 139 loci remained genome-wide significant at $P$ < 5E-8 in this subset. Among these 43 loci, 34 were still significant after conditional on T1D phenotype. For those loci that were not significant conditioning on the T1D phenotype, the differences in $P$-value were mostly marginal except for two SNPs in MHC region: rs1063355 located in *HLA-DQB1*, $P_{\text{unconditional}}$=1.1E-15, $P_{\text{conditional}}$=1.3E-4; rs2071479 located in *HLA-DOB*, $P_{\text{unconditional}}$=5.2E-7, $P_{\text{conditional}}$=5.8E-1, indicating that most T2D loci detected in this study were independent of T1D.

We could not find any publicly available data for LADA to perform the analyses above. However, a recent study[26] commented that "type 1 diabetes genetic risk score provides a reasonable approximation for all autoimmune diabetes". Under this hypothesis, our conclusion on T1D misdiagnosis is expected to hold for LADA. In addition, previous studies identified three T2D associations due to misdiagnosed LADA cases, i.e., *HLA-DQB1*[27,28], *INS*[27] and *PTPN22*[27]. Two of them (*INS* and *PTPN22*) were not genome-wide significant and *HLA-DQB1* was not a novel locus in our analysis.

In conclusion, the novel loci identified in this study were very unlikely due to misdiagnosed T1D or LADA cases.

**Supplementary Note 10 Acknowledgements**

**UKB:** This study has been conducted using UK Biobank resource under Application Number 12514. UK Biobank was established by the Wellcome Trust medical charity, Medical Research Council, Department of Health, Scottish Government and the Northwest Regional Development Agency. It has also had funding from the Welsh Assembly Government, British Heart Foundation and Diabetes UK.

**GERA** (dbGaP accession: phs000674.v2.p2)**:** The Genetic Epidemiology Research on Adult Health and Aging study was supported by grant RC2 AG036607 from the National Institutes of Health, grants from the Robert Wood Johnson Foundation, the Ellison Medical Foundation, the Wayne and Gladys Valley Foundation and Kaiser Permanente. The authors thank the Kaiser Permanente Medical Care Plan, Northern California Region (KPNC) members who have generously agreed to participate in the Kaiser Permanente Research Program on Genes, Environment and Health (RPGEH).

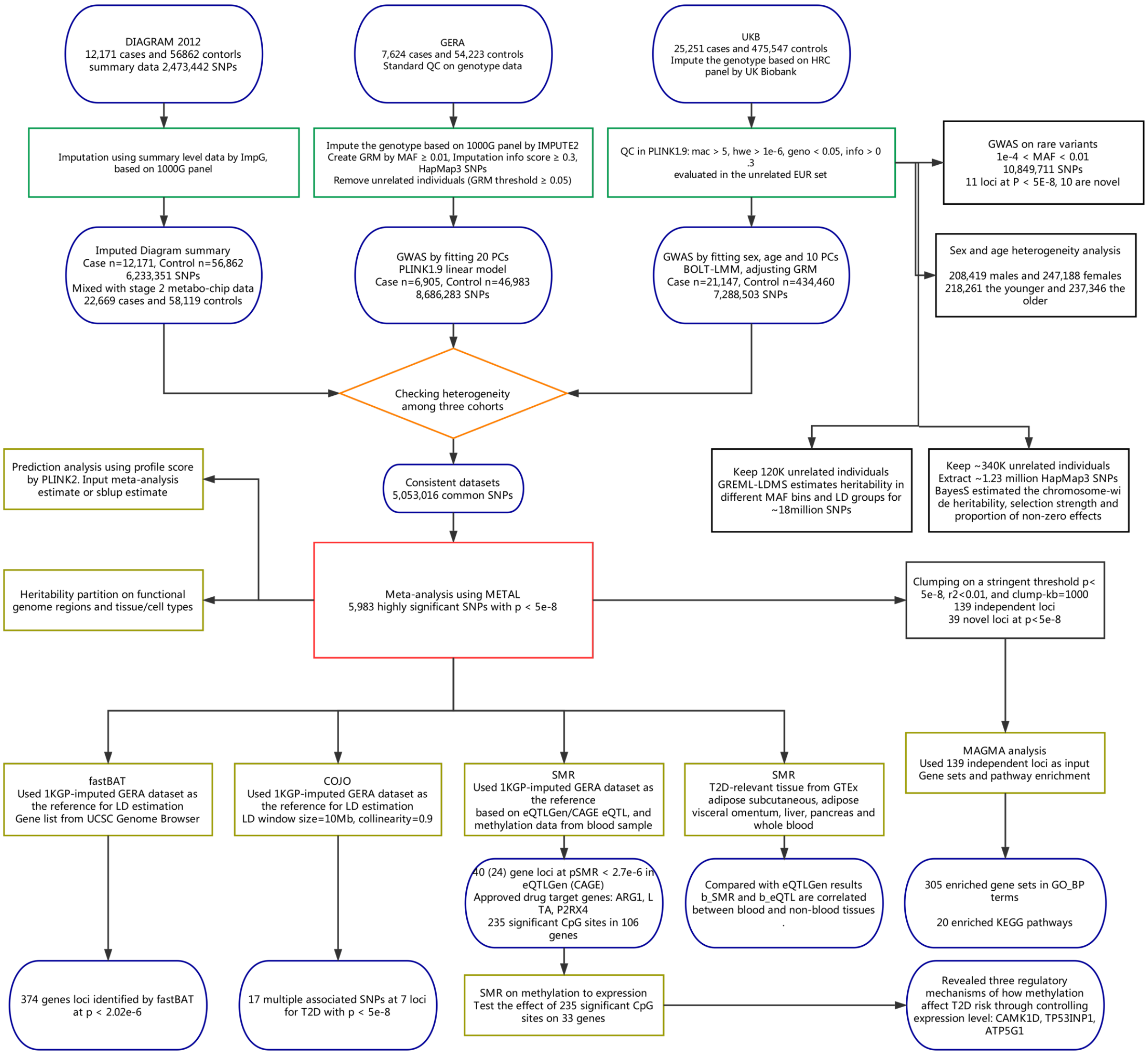


**Supplementary Figure 1** Schematic diagram of this study.

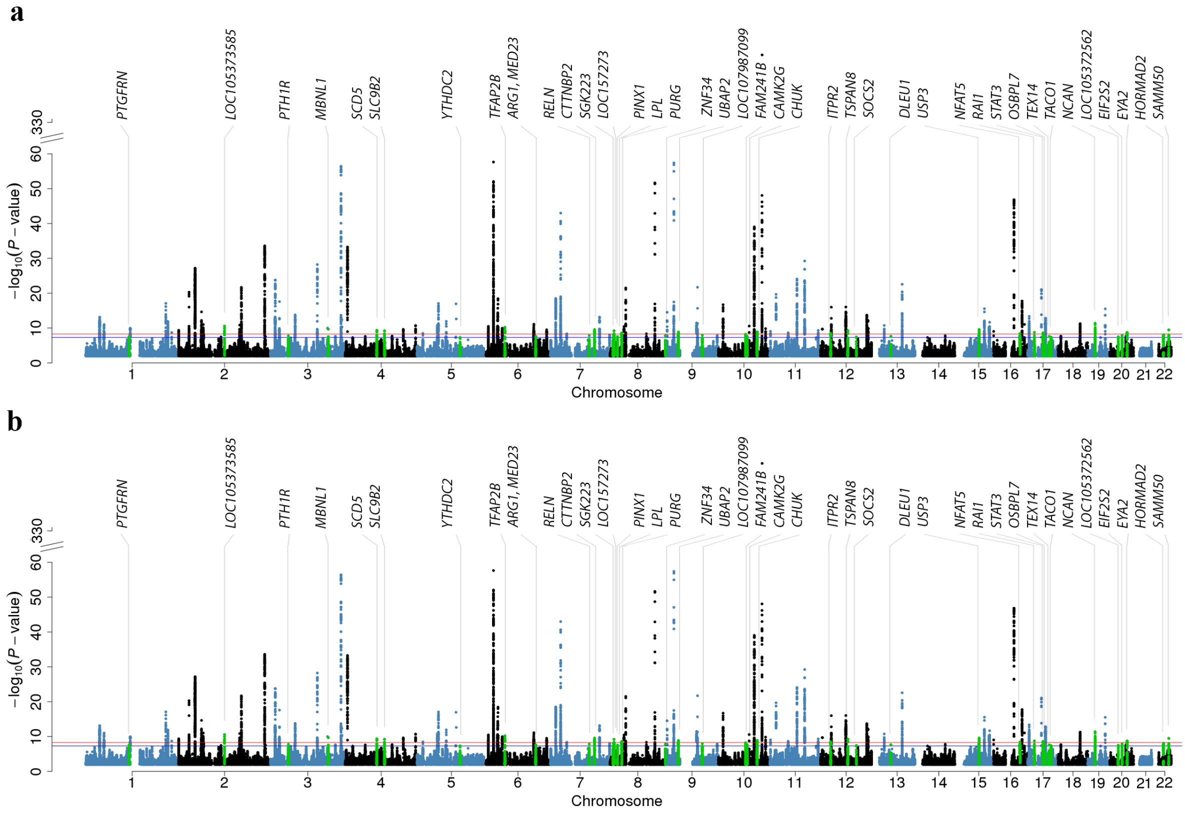


**Supplementary Figure 2** Manhattan plots of meta-analysis with the GERA cohort imputed to different imputation reference panels. a) GERA was imputed to the 1000 Genomes Project (1000G). b) GERA was imputed to the Haplotype Reference Consortium (HRC) using the Sanger imputation server (https://imputation.sanger.ac.uk/). Shown are the associations for ~5 million common variants.

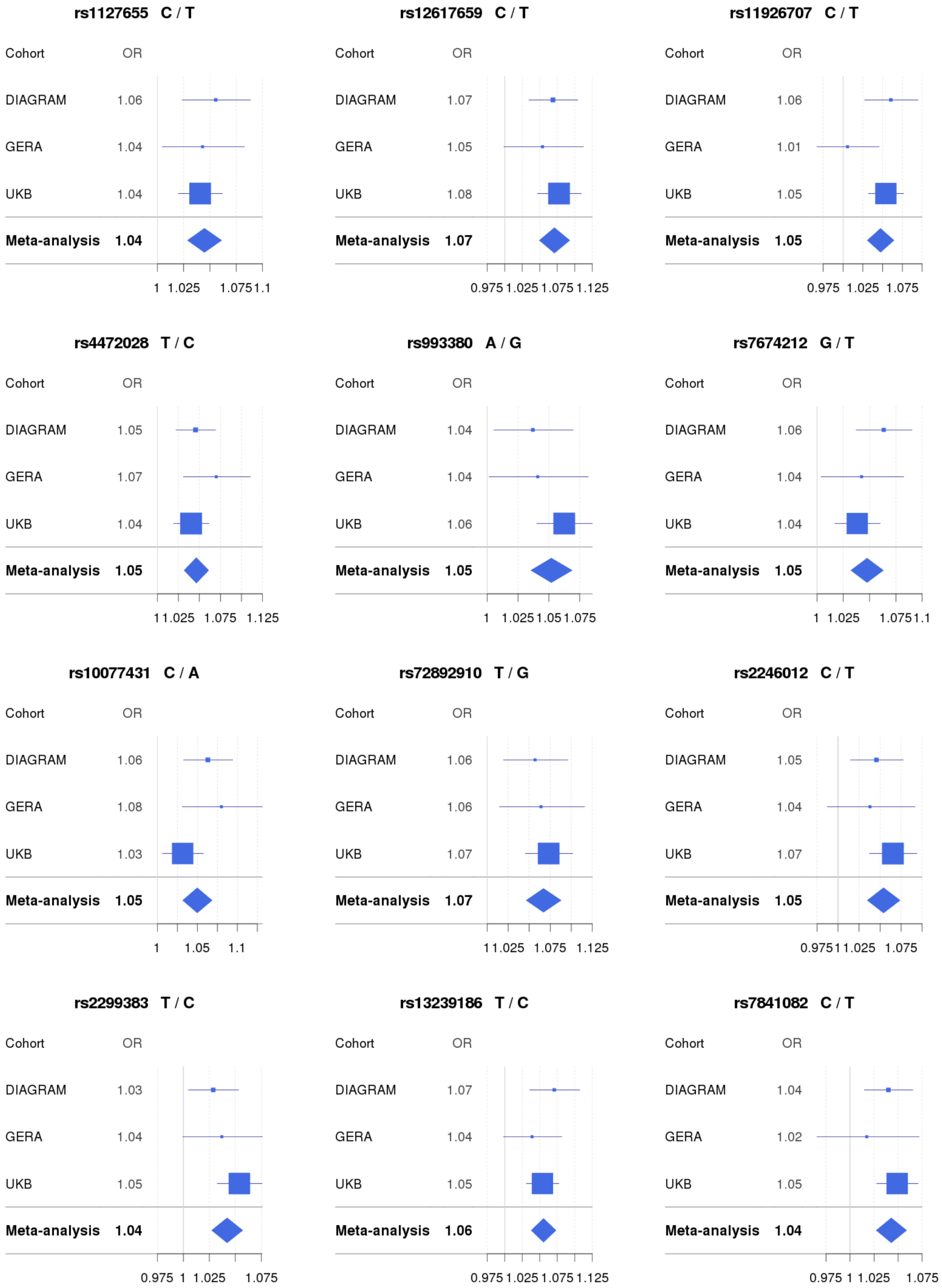
rs1127655 C / T
Cohort OR
DIAGRAM 1.06
GERA 1.04
UKB 1.04
Meta-analysis 1.04
1 1.025 1.075 1.1
rs12617659 C / T
Cohort OR
DIAGRAM 1.07
GERA 1.05
UKB 1.08
Meta-analysis 1.07
0.975 1.025 1.075 1.125
rs11926707 C / T
Cohort OR
DIAGRAM 1.06
GERA 1.01
UKB 1.05
Meta-analysis 1.05
0.975 1.025 1.075
rs4472028 T / C
Cohort OR
DIAGRAM 1.05
GERA 1.07
UKB 1.04
Meta-analysis 1.05
1 1.025 1.075 1.125
rs993380 A / G
Cohort OR
DIAGRAM 1.04
GERA 1.04
UKB 1.06
Meta-analysis 1.05
1 1.025 1.05 1.075
rs7674212 G / T
Cohort OR
DIAGRAM 1.06
GERA 1.04
UKB 1.04
Meta-analysis 1.05
1 1.025 1.075 1.1
rs10077431 C / A
Cohort OR
DIAGRAM 1.06
GERA 1.08
UKB 1.03
Meta-analysis 1.05
1 1.05 1.1
rs72892910 T / G
Cohort OR
DIAGRAM 1.06
GERA 1.06
UKB 1.07
Meta-analysis 1.07
1 1.025 1.075 1.125
rs2246012 C / T
Cohort OR
DIAGRAM 1.05
GERA 1.04
UKB 1.07
Meta-analysis 1.05
0.975 1 1.025 1.075
rs2299383 T / C
Cohort OR
DIAGRAM 1.03
GERA 1.04
UKB 1.05
Meta-analysis 1.04
0.975 1 1.025 1.075
rs13239186 T / C
Cohort OR
DIAGRAM 1.07
GERA 1.04
UKB 1.05
Meta-analysis 1.06
0.975 1.025 1.075 1.125
rs7841082 C / T
Cohort OR
DIAGRAM 1.04
GERA 1.02
UKB 1.05
Meta-analysis 1.04
0.975 1 1.025 1.075

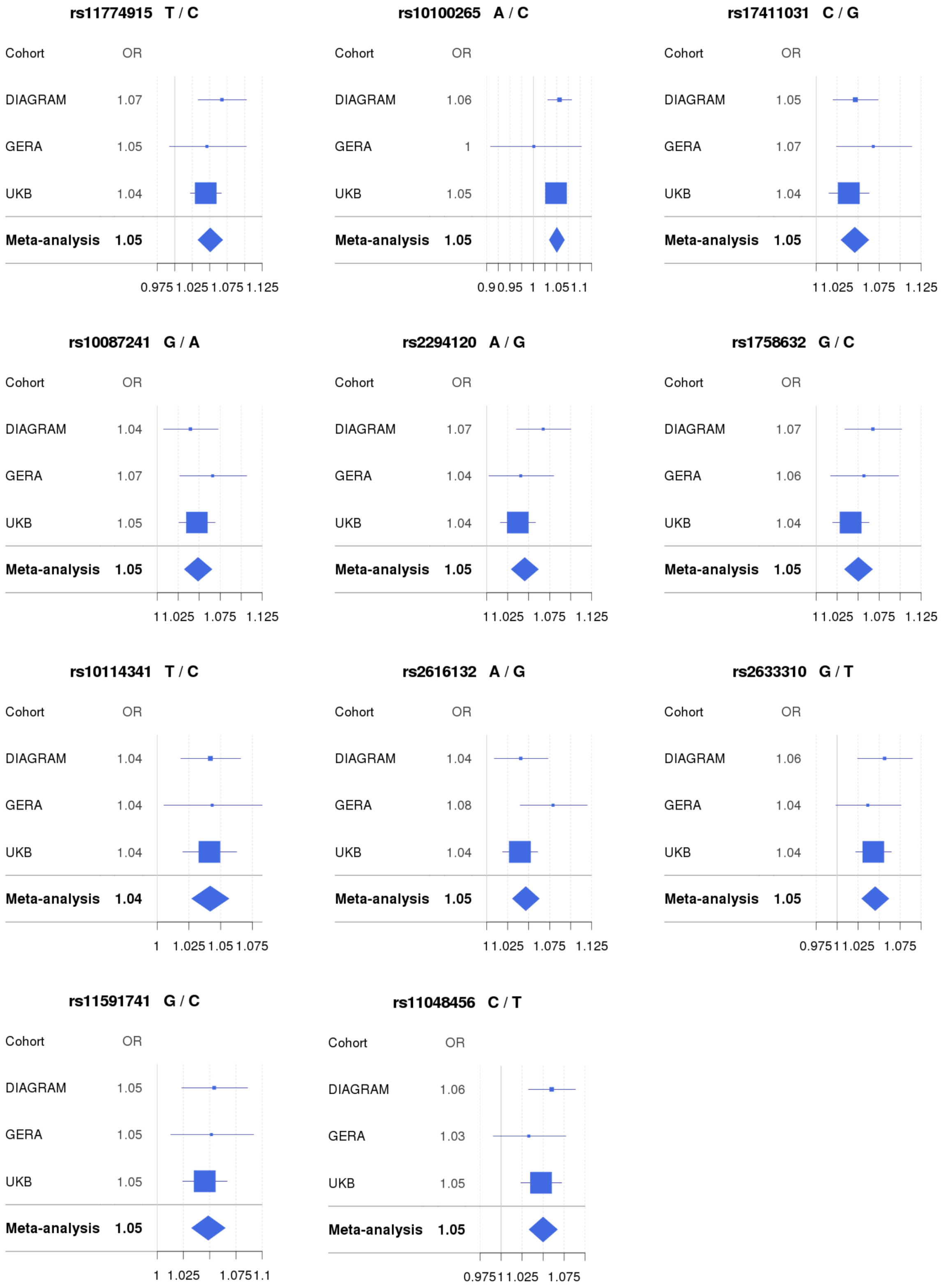
rs11774915 T / C
Cohort OR
DIAGRAM 1.07
GERA 1.05
UKB 1.04
Meta-analysis 1.05
0.975 1.025 1.075 1.125
rs10100265 A / C
Cohort OR
DIAGRAM 1.06
GERA 1
UKB 1.05
Meta-analysis 1.05
0.9 0.95 1 1.05 1.1
rs17411031 C / G
Cohort OR
DIAGRAM 1.05
GERA 1.07
UKB 1.04
Meta-analysis 1.05
1 1.025 1.075 1.125
rs10087241 G / A
Cohort OR
DIAGRAM 1.04
GERA 1.07
UKB 1.05
Meta-analysis 1.05
1 1.025 1.075 1.125
rs2294120 A / G
Cohort OR
DIAGRAM 1.07
GERA 1.04
UKB 1.04
Meta-analysis 1.05
1 1.025 1.075 1.125
rs1758632 G / C
Cohort OR
DIAGRAM 1.07
GERA 1.06
UKB 1.04
Meta-analysis 1.05
1 1.025 1.075 1.125
rs10114341 T / C
Cohort OR
DIAGRAM 1.04
GERA 1.04
UKB 1.04
Meta-analysis 1.04
1 1.025 1.05 1.075
rs2616132 A / G
Cohort OR
DIAGRAM 1.04
GERA 1.08
UKB 1.04
Meta-analysis 1.05
1 1.025 1.075 1.125
rs2633310 G / T
Cohort OR
DIAGRAM 1.06
GERA 1.04
UKB 1.04
Meta-analysis 1.05
0.975 1 1.025 1.075
rs11591741 G / C
Cohort OR
DIAGRAM 1.05
GERA 1.05
UKB 1.05
Meta-analysis 1.05
1 1.025 1.075 1.1
rs11048456 C / T
Cohort OR
DIAGRAM 1.06
GERA 1.03
UKB 1.05
Meta-analysis 1.05
0.975 1 1.025 1.075

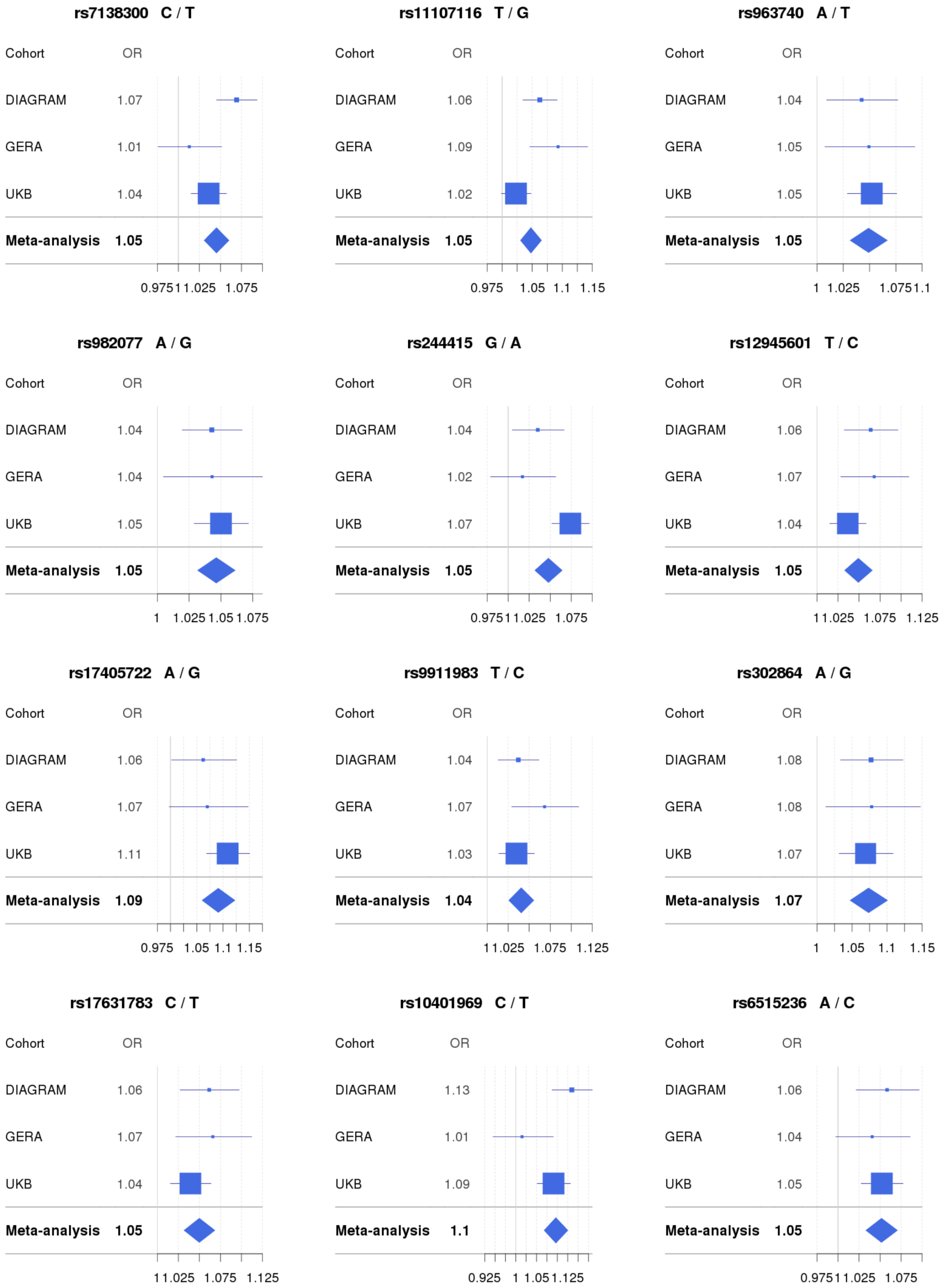
rs7138300 C / T
Cohort OR
DIAGRAM 1.07
GERA 1.01
UKB 1.04
Meta-analysis 1.05
0.975 1 1.025 1.075
rs11107116 T / G
Cohort OR
DIAGRAM 1.06
GERA 1.09
UKB 1.02
Meta-analysis 1.05
0.975 1.05 1.1 1.15
rs963740 A / T
Cohort OR
DIAGRAM 1.04
GERA 1.05
UKB 1.05
Meta-analysis 1.05
1 1.025 1.075 1.1
rs982077 A / G
Cohort OR
DIAGRAM 1.04
GERA 1.04
UKB 1.05
Meta-analysis 1.05
1 1.025 1.05 1.075
rs244415 G / A
Cohort OR
DIAGRAM 1.04
GERA 1.02
UKB 1.07
Meta-analysis 1.05
0.975 1 1.025 1.075
rs12945601 T / C
Cohort OR
DIAGRAM 1.06
GERA 1.07
UKB 1.04
Meta-analysis 1.05
1 1.025 1.075 1.125
rs17405722 A / G
Cohort OR
DIAGRAM 1.06
GERA 1.07
UKB 1.11
Meta-analysis 1.09
0.975 1.05 1.1 1.15
rs9911983 T / C
Cohort OR
DIAGRAM 1.04
GERA 1.07
UKB 1.03
Meta-analysis 1.04
1 1.025 1.075 1.125
rs302864 A / G
Cohort OR
DIAGRAM 1.08
GERA 1.08
UKB 1.07
Meta-analysis 1.07
1 1.05 1.1 1.15
rs17631783 C / T
Cohort OR
DIAGRAM 1.06
GERA 1.07
UKB 1.04
Meta-analysis 1.05
1 1.025 1.075 1.125
rs10401969 C / T
Cohort OR
DIAGRAM 1.13
GERA 1.01
UKB 1.09
Meta-analysis 1.1
0.925 1 1.05 1.125
rs6515236 A / C
Cohort OR
DIAGRAM 1.06
GERA 1.04
UKB 1.05
Meta-analysis 1.05
0.975 1 1.025 1.075

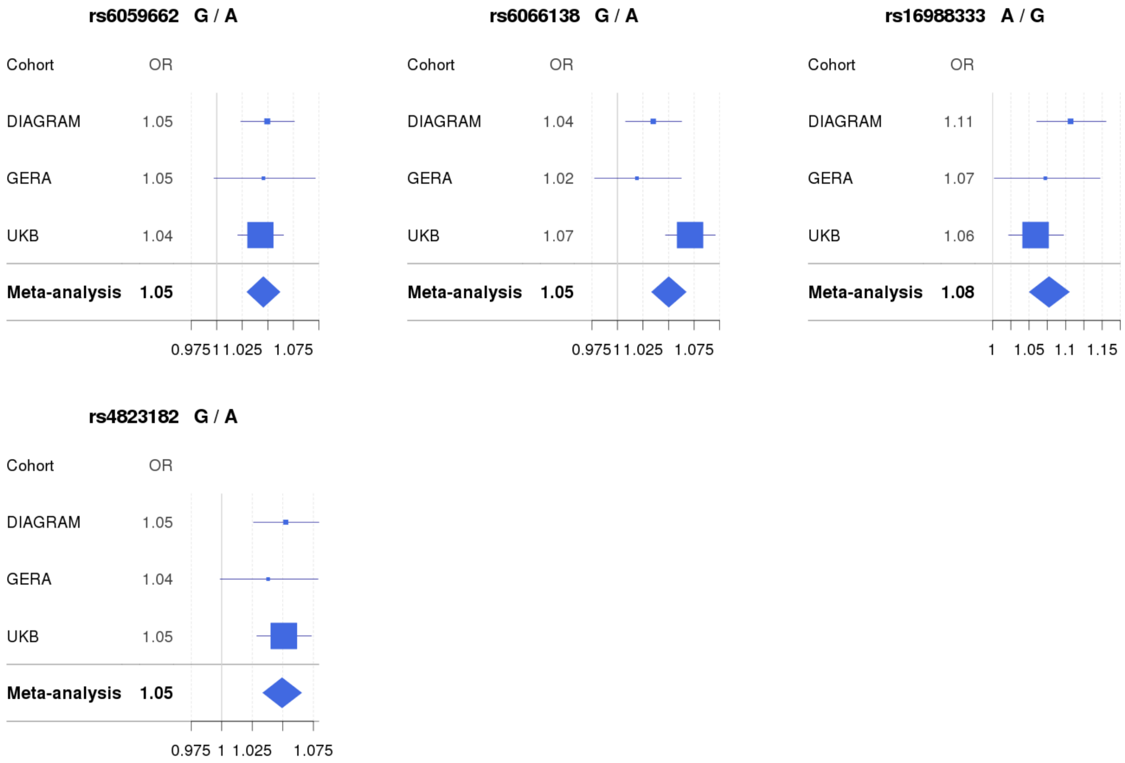


**Supplementary Figure 3** Forest plots of the 39 novel loci associated with T2D at $P < 5E\text{-}8$. Error bars represent the 95% confidence intervals. Area of the square (or rhombus) denotes the sample size.

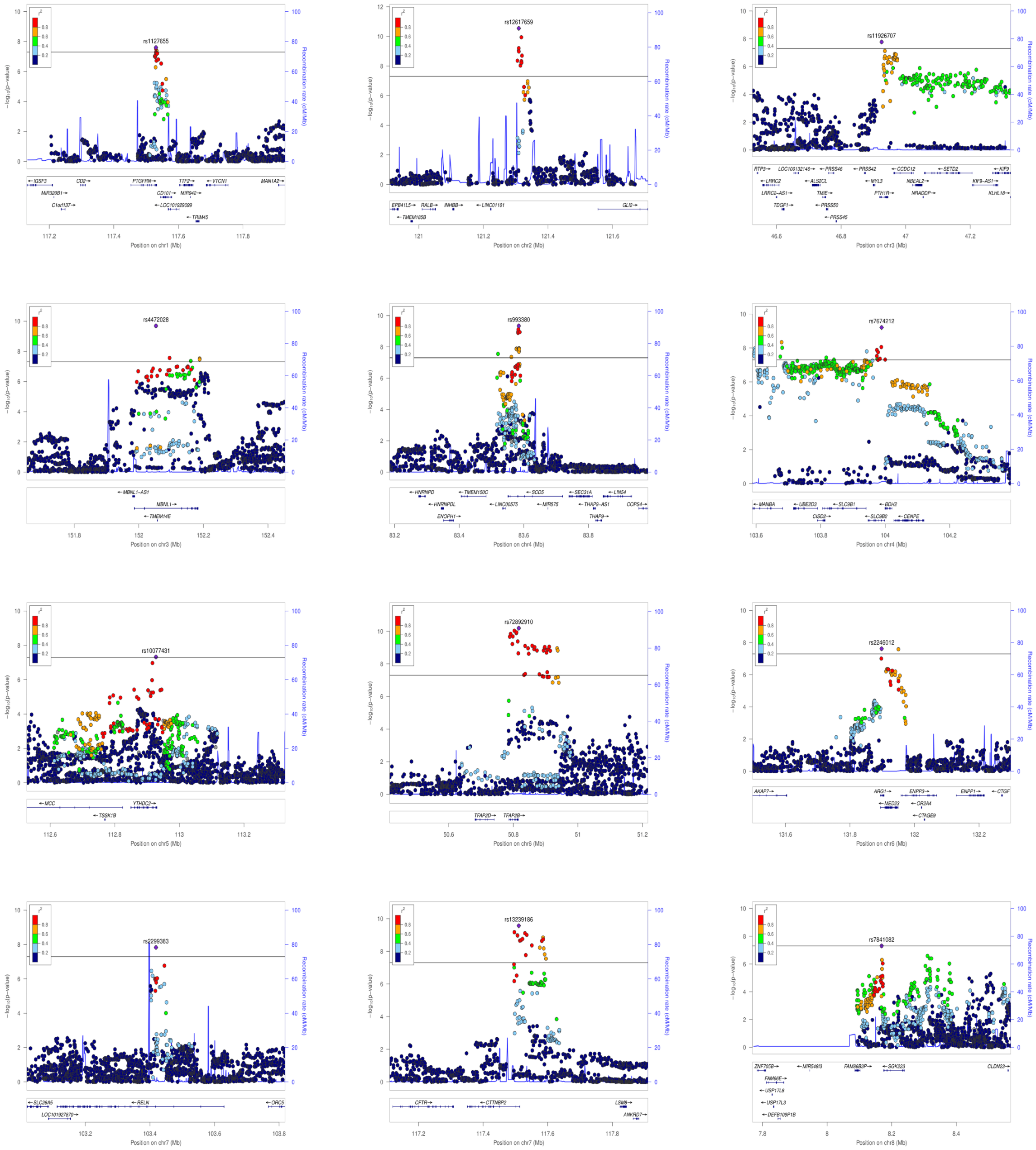
rs1127655
Position on chr1 (Mb)
rs12617659
Position on chr2 (Mb)
rs11926707
Position on chr3 (Mb)
rs4472028
Position on chr3 (Mb)
rs993380
Position on chr4 (Mb)
rs7674212
Position on chr4 (Mb)
rs10077431
Position on chr5 (Mb)
rs72892910
Position on chr6 (Mb)
rs2246012
Position on chr6 (Mb)
rs2299383
Position on chr7 (Mb)
rs13239186
Position on chr7 (Mb)
rs7841082
Position on chr8 (Mb)
Recombination rate (cM/Mb)

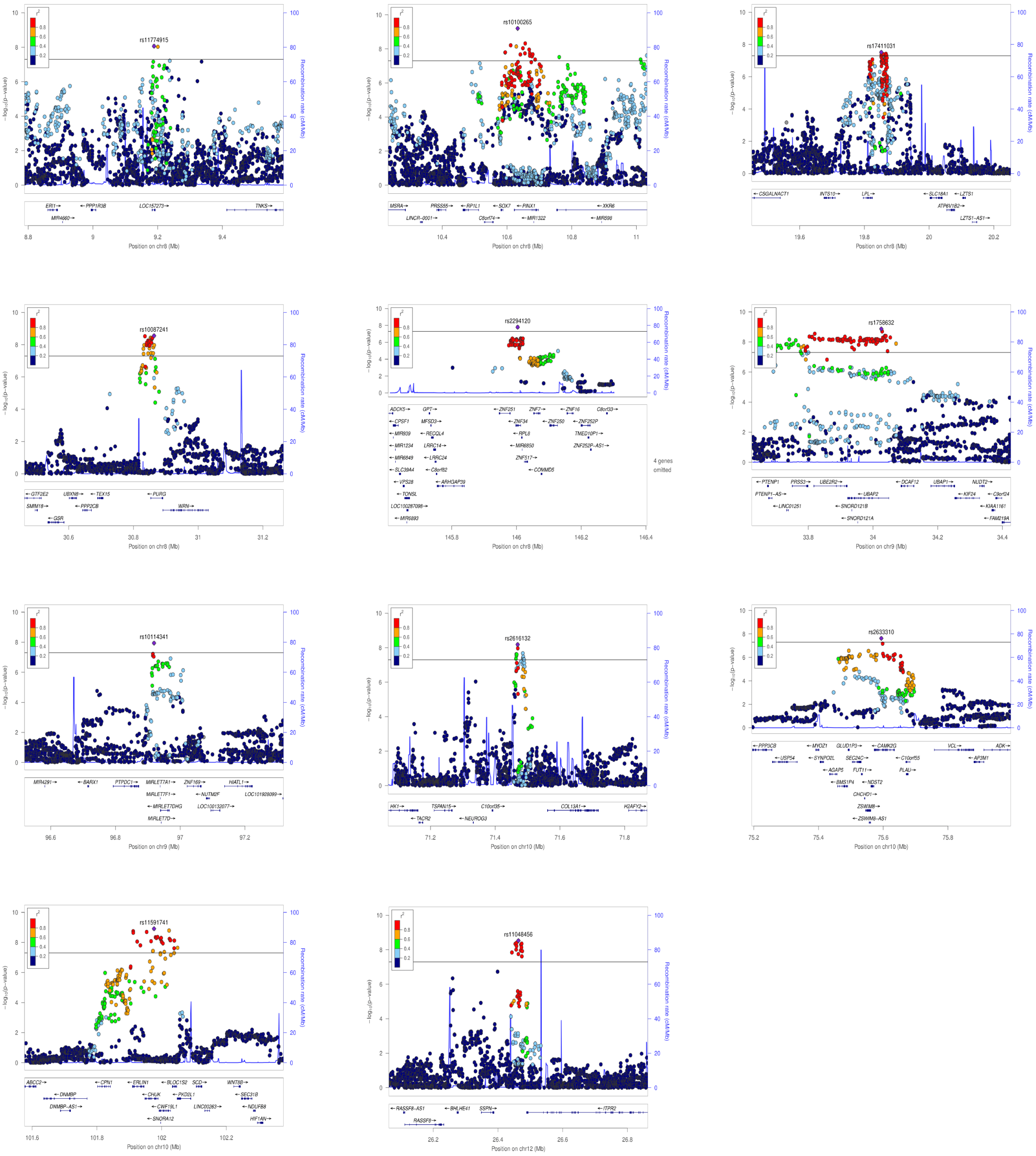

rs11774915
Position on chr8 (Mb)
rs10100265
Position on chr8 (Mb)
rs17411031
Position on chr8 (Mb)
rs10087241
Position on chr8 (Mb)
rs2294120
Position on chr8 (Mb)
rs1758632
Position on chr9 (Mb)
rs10114341
Position on chr9 (Mb)
rs2616132
Position on chr10 (Mb)
rs2633310
Position on chr10 (Mb)
rs11591741
Position on chr10 (Mb)
rs11048456
Position on chr12 (Mb)

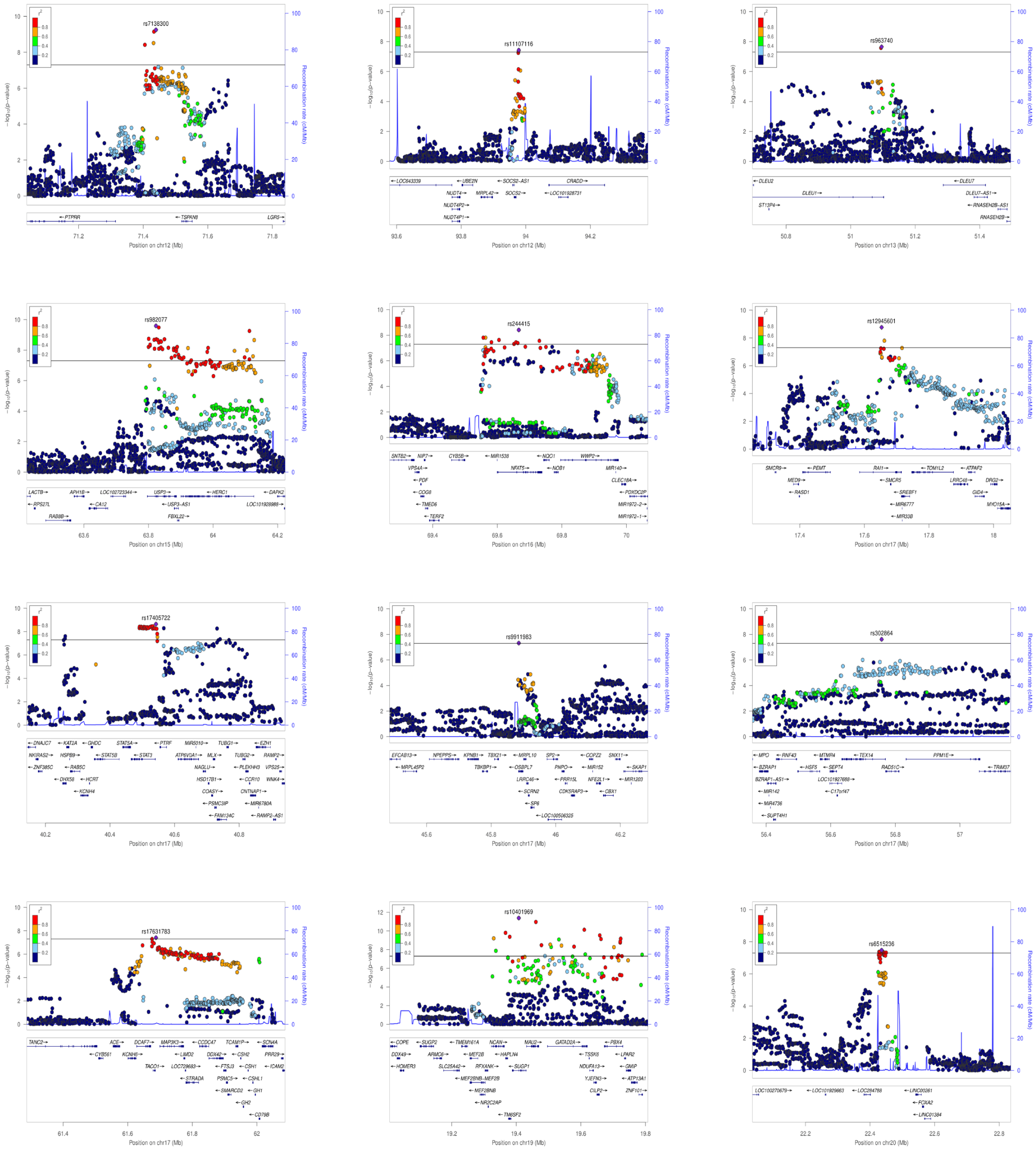
rs7138300
Position on chr12 (Mb)
rs11107116
Position on chr12 (Mb)
rs963740
Position on chr13 (Mb)
rs982077
Position on chr15 (Mb)
rs244415
Position on chr16 (Mb)
rs12945601
Position on chr17 (Mb)
rs17405722
Position on chr17 (Mb)
rs9911983
Position on chr17 (Mb)
rs302864
Position on chr17 (Mb)
rs17631783
Position on chr17 (Mb)
rs10401969
Position on chr19 (Mb)
rs6515236
Position on chr20 (Mb)

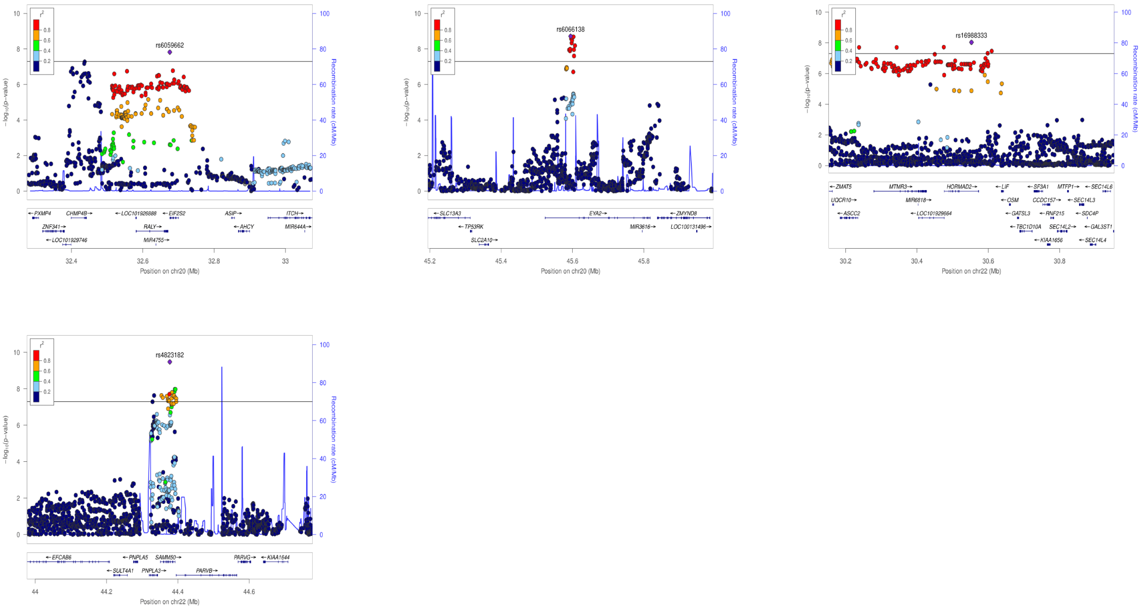


**Supplementary Figure 4** Regional plots of the 39 novel loci identified from the meta-analysis at $P < 5 \times 10^{-8}$. Each point represents a SNP passing quality control in the meta-analysis plotted with its $P$ value (on a -log10 scale) against its genomic position (NCBI Build 37). In each plot, the lead SNP is represented by the purple symbol. The color coding of all other SNPs indicate LD with the lead SNP (estimated by CEU $r^2$ values from Phase I 1000 Genomes): red, $r^2 \geqslant 0.8$; gold, $0.6 \leqslant r^2 < 0.8$; green, $0.4 \leqslant r^2 < 0.6$; cyan, $0.2 \leqslant r^2 < 0.4$; blue, $r^2 < 0.2$; gray, $r^2$ unknown. Recombination rates are estimated from Phase I 1000 Genomes, and gene annotations are taken from the UCSC genome browser.

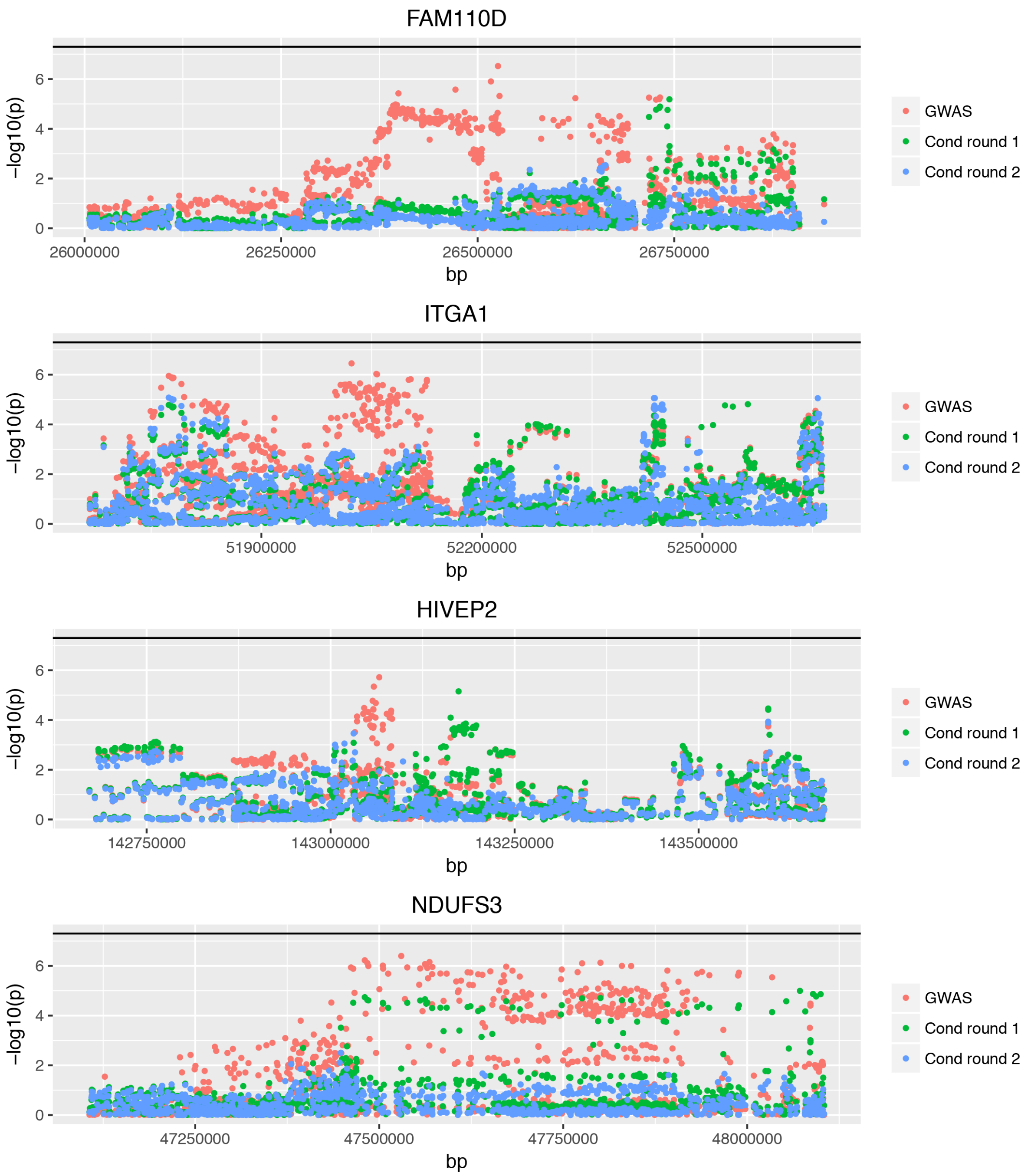


**Supplementary Figure 5** Conditional association analysis at the four novel loci identified by the GCTA-fastBAT analysis.

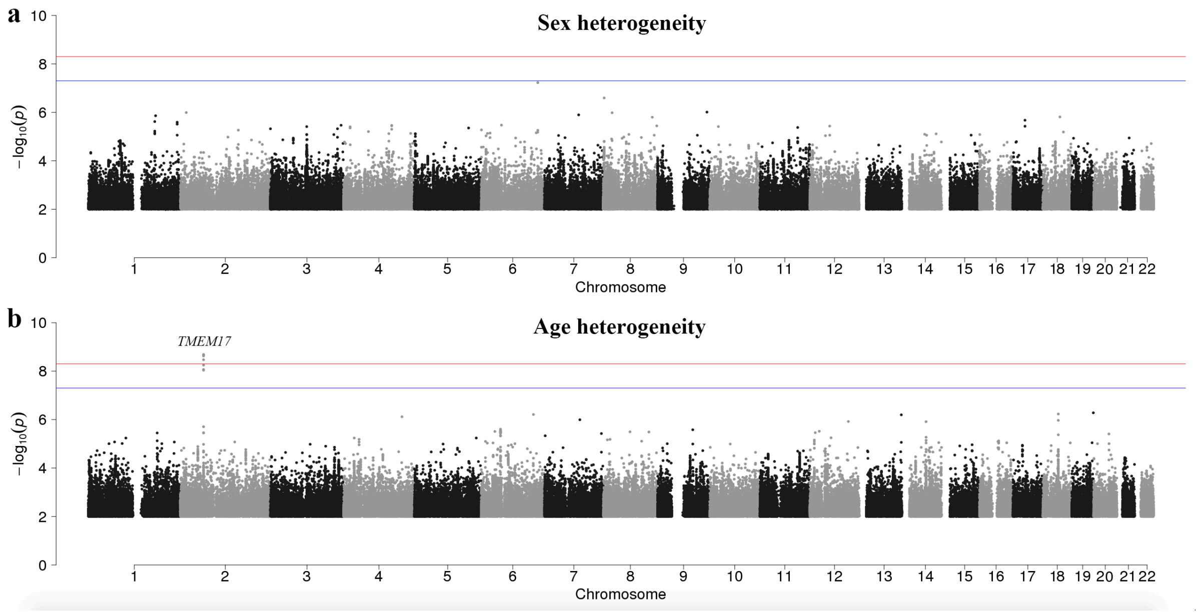


**Supplementary Figure 6** Heterogeneity in SNP effect between sex (or age) groups in UKB. Shown are the Manhattan plots from the heterogeneity tests between sex (or age) groups for all 18,138,214 variants (including the rare variants) (**Supplementary Note**). The x-axis is the chromosome number and the y-axis is the –log10 of heterogeneity *P*-value. The blue lines represent a genome-wide significance level at $P < 5\times10^{-8}$, and the red lines represent a threshold of 5E-9 (as suggested by Wu *et al.*[29] for GWAS with both common and rare variants from imputation). SNPs with $P_{\text{heter}} > 0.01$ are omitted.

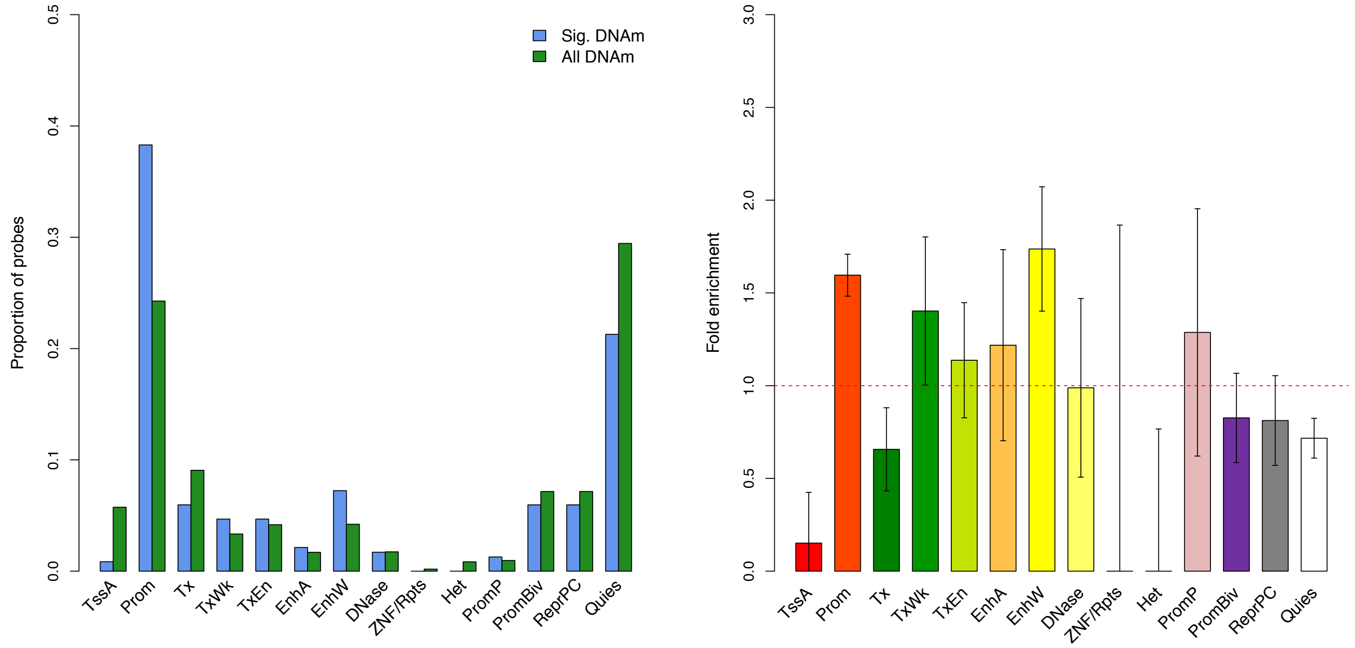


**Supplementary Figure 7** Enrichment of the 235 T2D-associated DNAm sites in functional categories. a) Distribution of the T2D-associated DNAm probes ("Sig. DNAm", blue) across the 14 functional categories in comparison with that of all DNAm probes in the data ("All DNAm", green). b) Fold enrichment: a comparison of the T2D-associated probes with the same number of probes sampled repeatedly at random with the variance of each probe matched. Error bar represents the standard error of an estimate obtained from 500 random samples. The 14 functional annotation categories are: TssA, active transcription start site; Prom, upstream/downstream TSS promoter; Tx, actively transcribed state; TxWk, weak transcription; TxEn, transcribed and regulatory Prom/Enh; EnhA, active enhancer; EnhW, weak enhancer; DNase, primary DNase; ZNF/Rpts, state associated with zinc finger protein genes; Het, constitutive heterochromatin; PromP, Poised promoter; PromBiv, bivalent regulatory states; ReprPC, repressed Polycomb states; and Quies, a quiescent state.

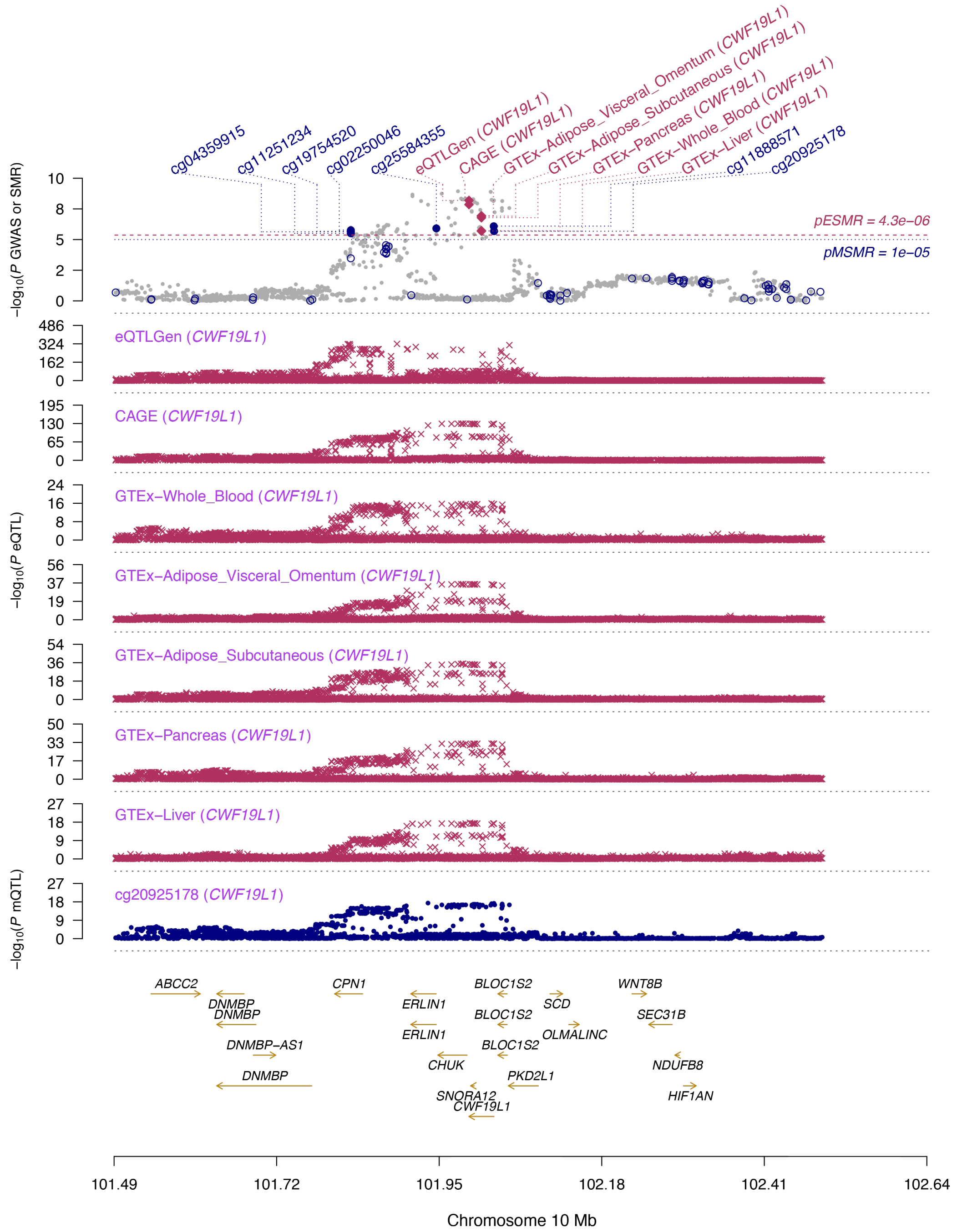


**Supplementary Figure 8** Association signals of the expression level of *CWF19L1* with the cis-SNPs in eQTLGen and CAGE in comparison with those in five GTEx tissues. Shown are the results from the SMR analysis that integrates data from GWAS, eQTL and mQTL studies. The top plot shows -$\log_{10}$($P$-value) from our GWAS meta-analysis. Red diamonds and blue circles represent -$\log_{10}$($P$-value) from the SMR tests for associations of gene expression and DNA methylation probes with T2D, respectively. Solid diamonds and circles represent the probes not rejected by the HEIDI test. The second plot shows -$\log_{10}$($P$-value) of SNP associations with gene expression probes (tagging *CWF19L1*) in eQTLGen, CAGE and five GTEx tissues. The third plot shows -$\log_{10}$($P$-value) of SNP associations with a DNA methylation probe cg20925178.

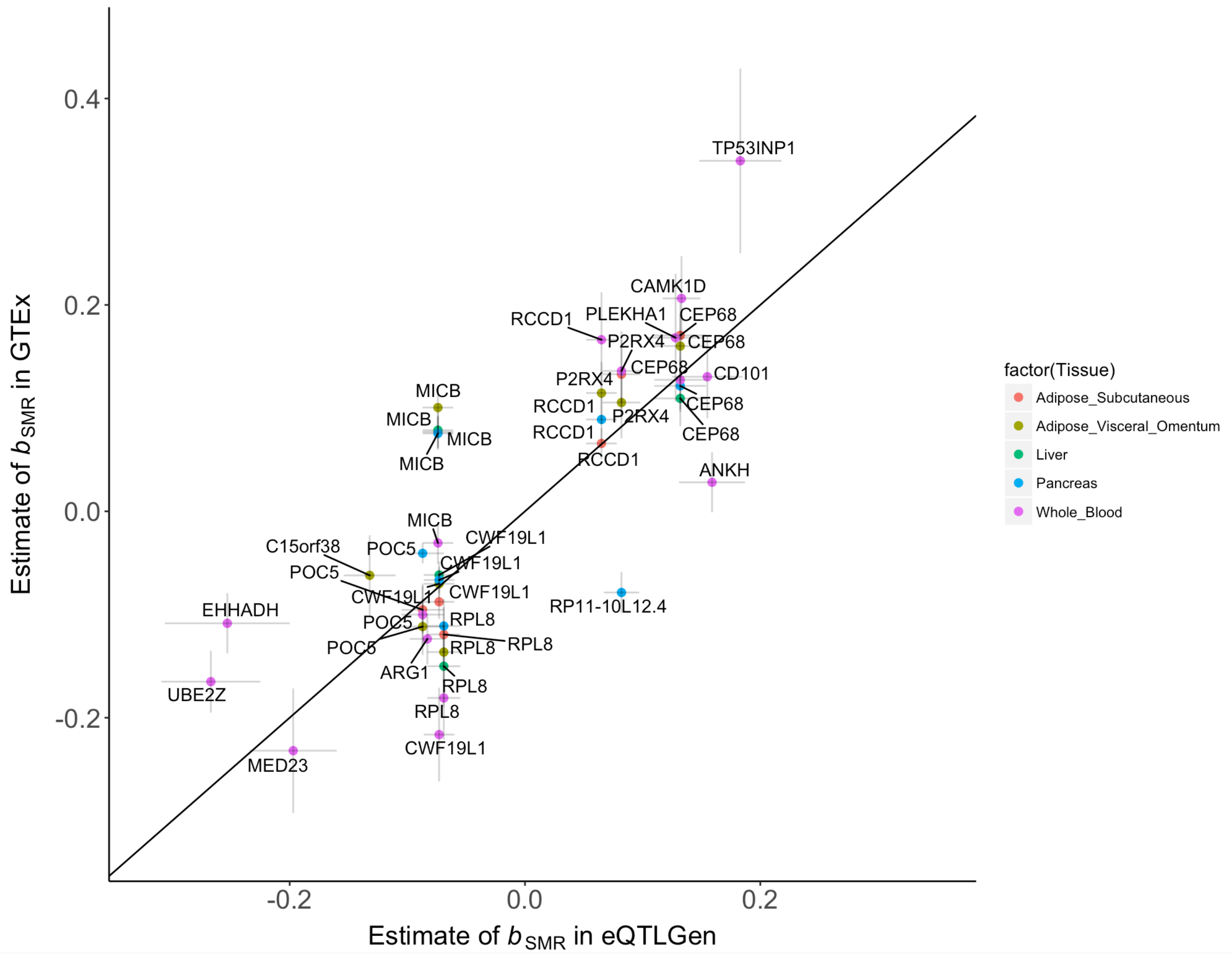


**Supplementary Figure 9** Estimates of SMR effects in eQTLGen vs. those in the five GTEx tissues. The SMR effect ($b_{SMR}$) is defined as the effect of the expression level of a gene on T2D risk[30]. Shown are the 18 genes detected using the eQTLGen data with $b_{SMR}$ estimated in eQTLGen plotted against those estimated in the GTEx tissues (including pancreas, liver and/or adiposes). The MHC region was not included in the analysis. Each dot represents a gene with colors indicating different tissues in GTEx. Error bar represents the standard error for an estimate of $b_{SMR}$. The correlation of $b_{SMR}$ estimates between eQTLGen and GTEx is 0.80.

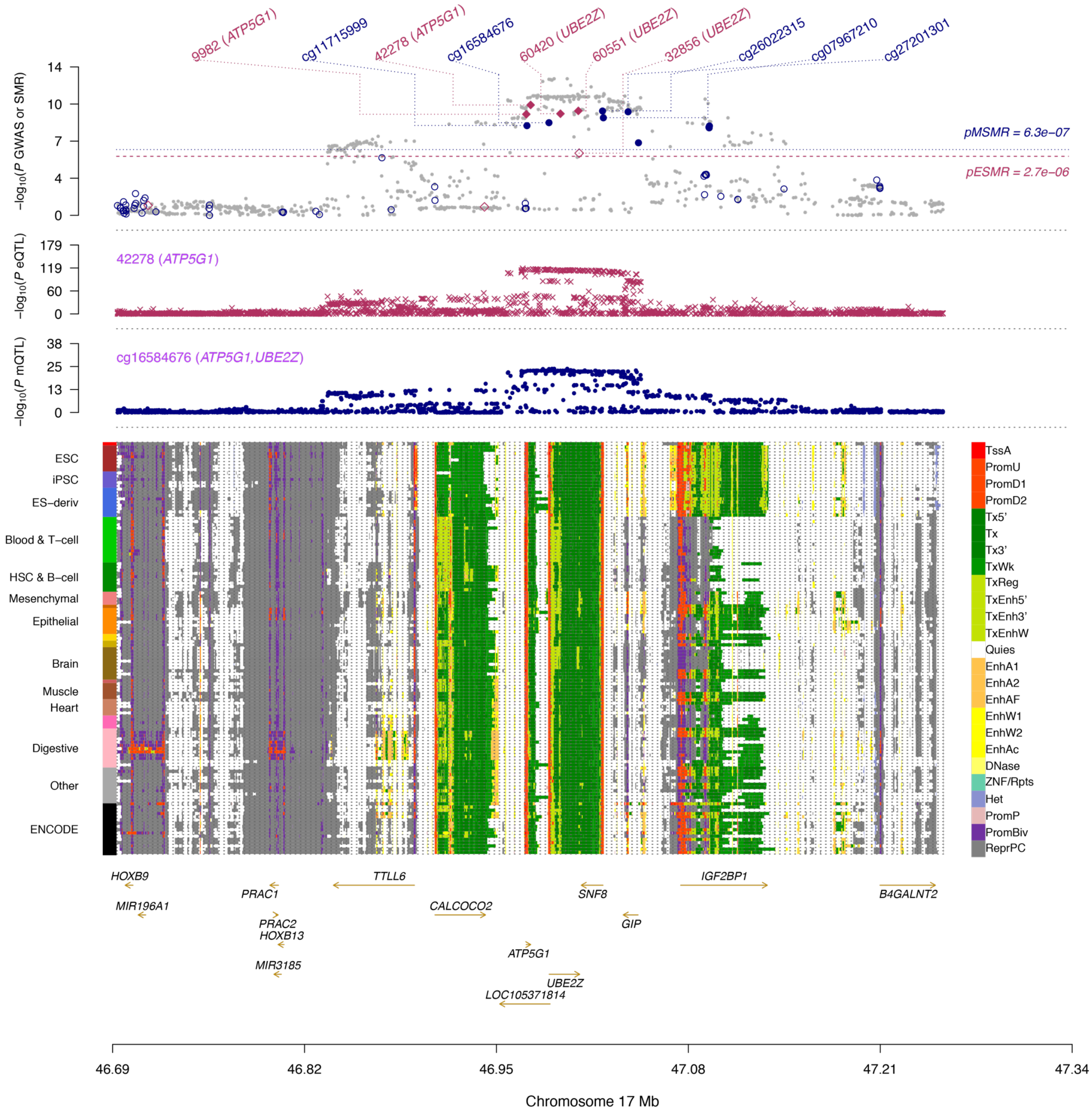


**Supplementary Figure 10a** Prioritizing genes and regulatory elements at the *ATP5G1* locus. Shown are the results from the SMR analysis that integrates data from GWAS, eQTL and mQTL studies. The top plot shows -log10(*P*-value) of SNPs from the GWAS meta-analysis for T2D. Red diamonds and blue circles represent -log10(*P*-value) from the SMR tests for associations of gene expression and DNAm probes with T2D, respectively. Solid diamonds and circles represent the probes not rejected by the HEIDI test. The second plot shows -log10(*P*-value) of the SNP association for gene expression probe 42278 (tagging *ATP5G1*). The third plot shows -log10(*P*-value) of the SNP association with DNAm probe cg16584676. The bottom plot shows 25 chromatin state annotations (indicated by colours) of 127 samples from Roadmap Epigenomics Mapping Consortium (REMC) for different primary cells and tissue types (rows).

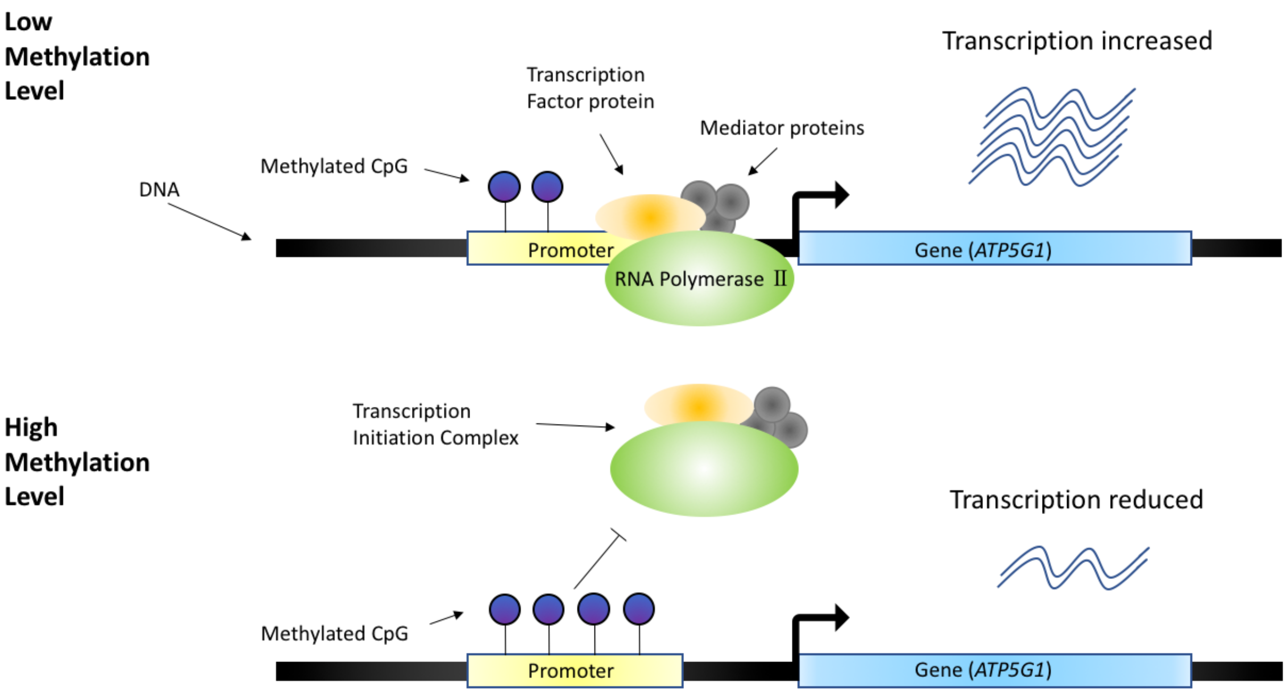


**Supplementary Figure 10b** Hypothesized mechanism of how DNA methylation affect the expression level of *ATP5G1*. When the methylation level of the promoter region is low, the RNA polymerase II binds to the promoter region with the assistance from transcription factors (TF), and initiate the transcription. However, if the promoter region is highly methylated, it would obstruct the binding of RNA polymerase II to promoter region, which leads to the reduction of transcription.

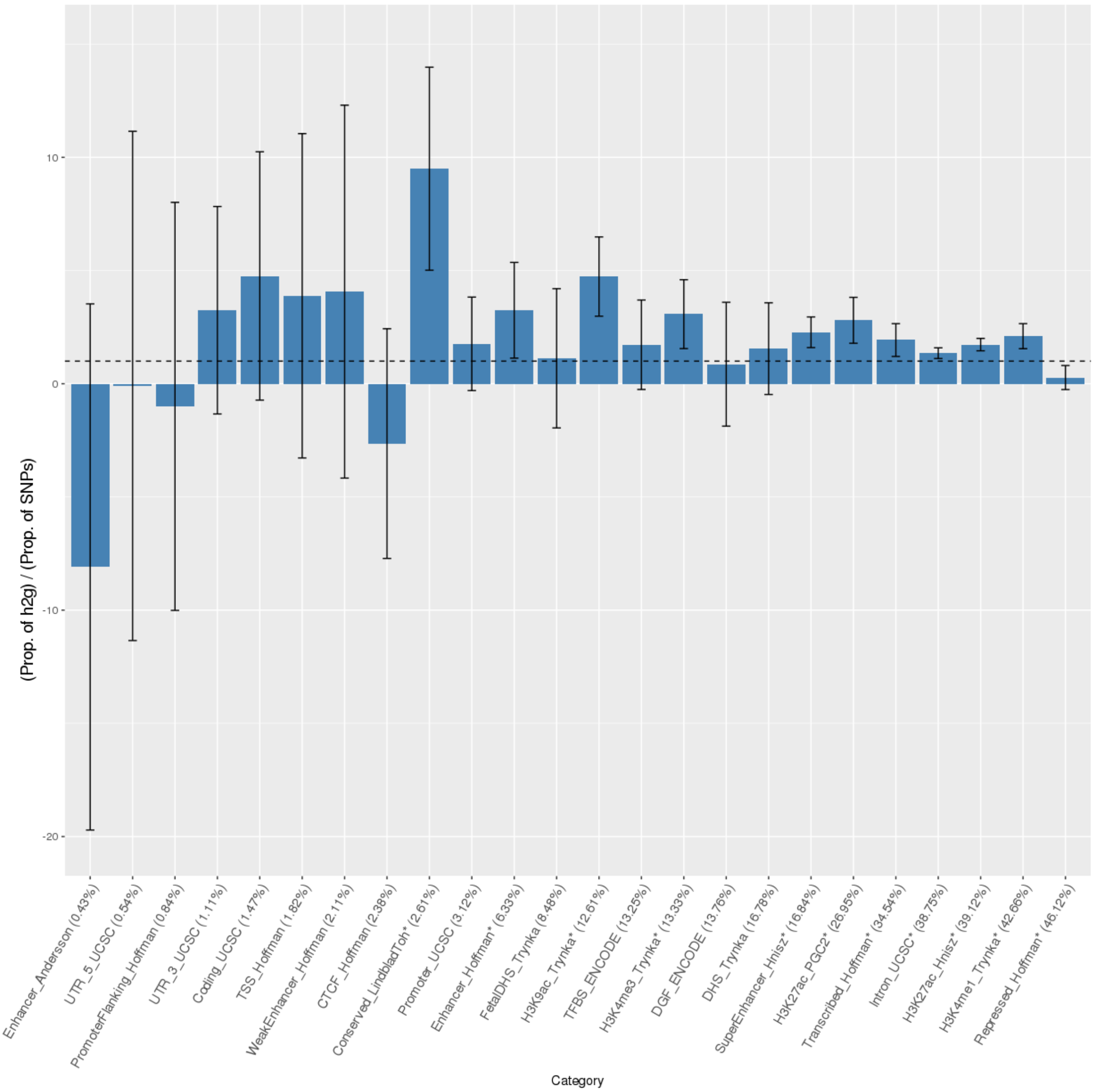


**Supplementary Figure 11** Enrichment of the variance explained by SNPs for T2D in 24 functional annotations. Shown are the results from the LD score regression based functional partitioning analysis[31]. The 24 functional annotations are defined in Finucane *et al.*[31]. Annotations are ordered by proportion of SNPs. Error bar represents the 95% confidential interval around the estimate of enrichment, and the asterisk indicates a significant estimate at $P < 0.05$ after Bonferroni correction for 24 tests. CTCF, CCCTC-binding factor. DGF, digital genomic footprint. DHS, DNase I hypersensitive site. TFBS, transcription factor binding site. TSS, transcription start site.

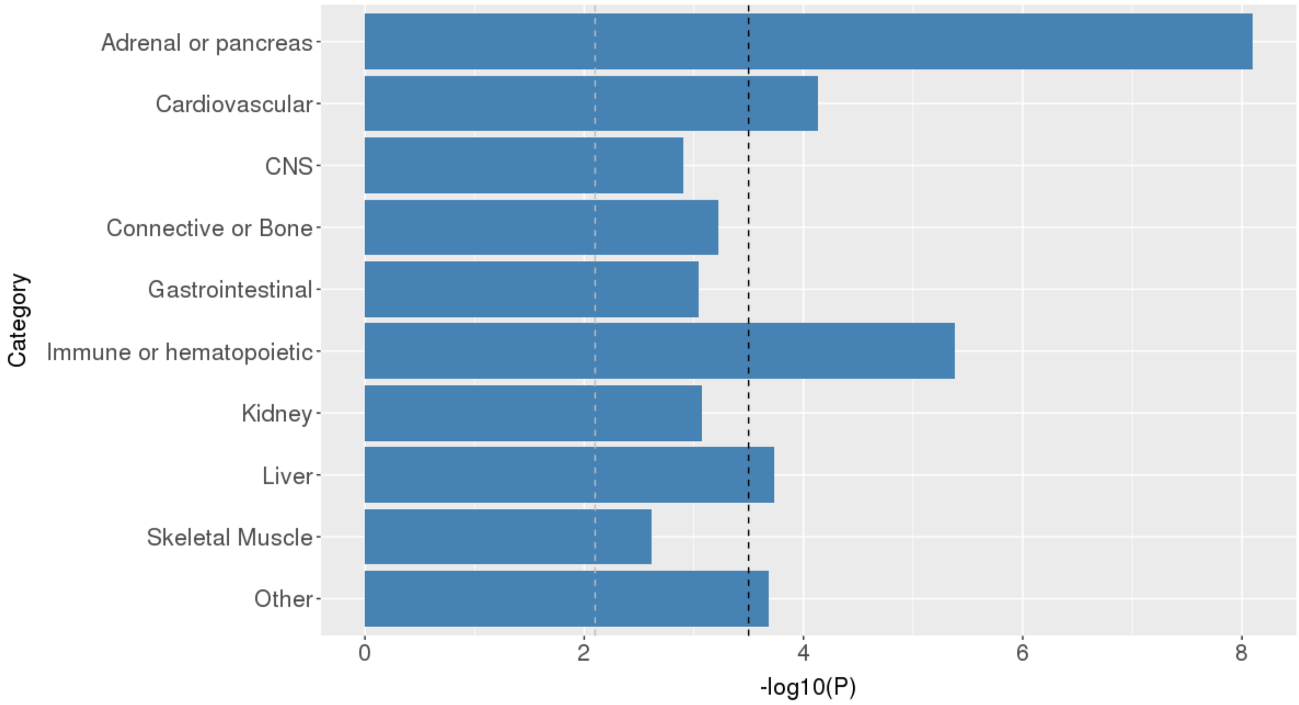


**Supplementary Figure 12** Enrichment of the variance explained by SNPs for T2D in 10 cell type groups. Shown are the results from the LD score regression based functional partitioning analysis[31]. The black dashed lines at -log10 ($P$) = 3.6 is the significance level after Bonferroni correction. The grey dashed lines at -log10 ($P$) = 2.3 is the threshold at FDR < 0.05.

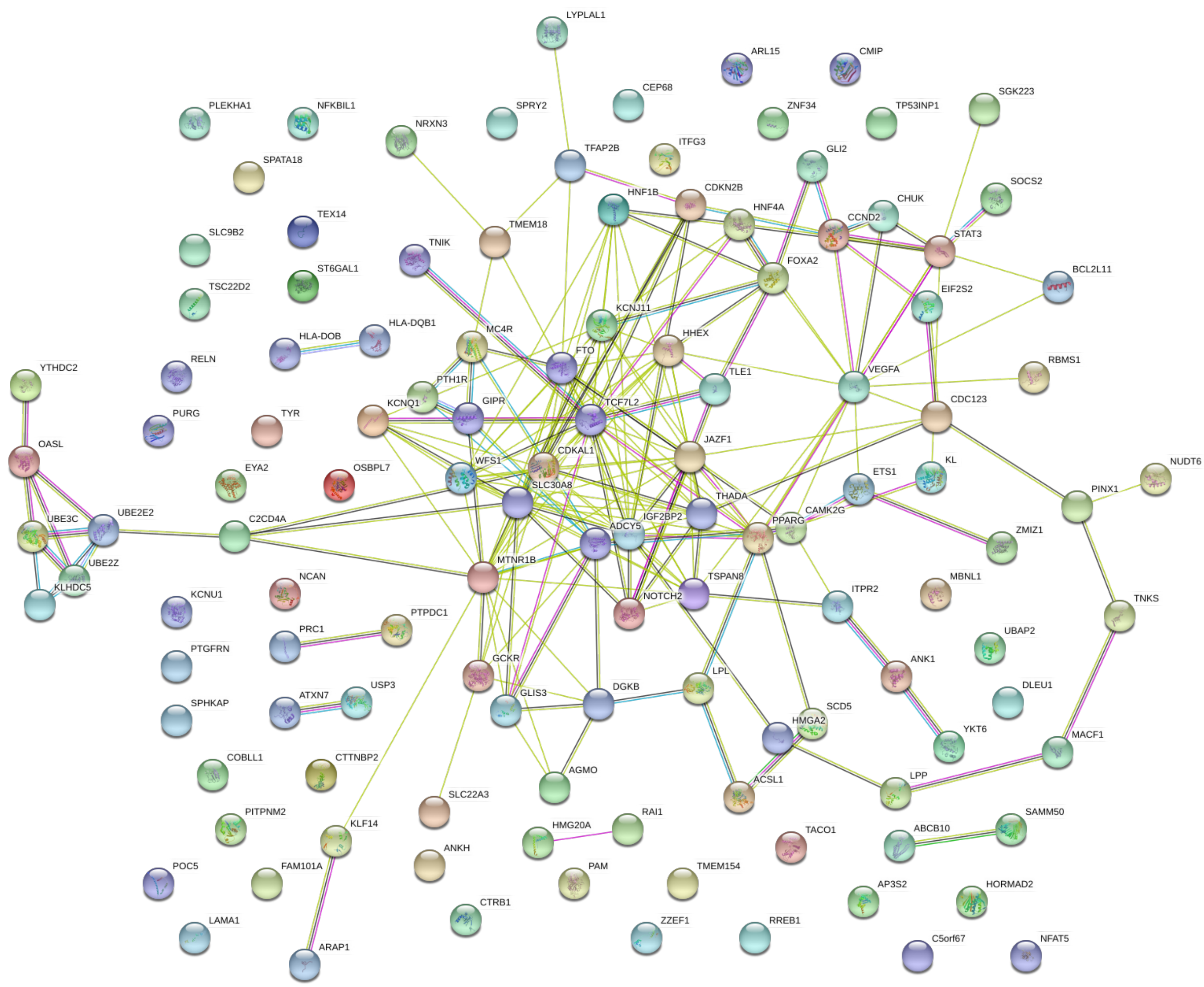


**Supplementary Figure 13** Enrichment of the T2D-associted genes in protein-protein interaction network. There are 123 nodes and 210 edges in total generated by STRING v10. Network nodes represent proteins. Small node represents protein of unknown 3D structure while large node represents that the 3D structure is known or predicted. Edges represent protein-protein associations. Dark purple edge represents experimentally determined interaction, light purple edge represents protein homology, and yellow edge represents interaction from text-mining.

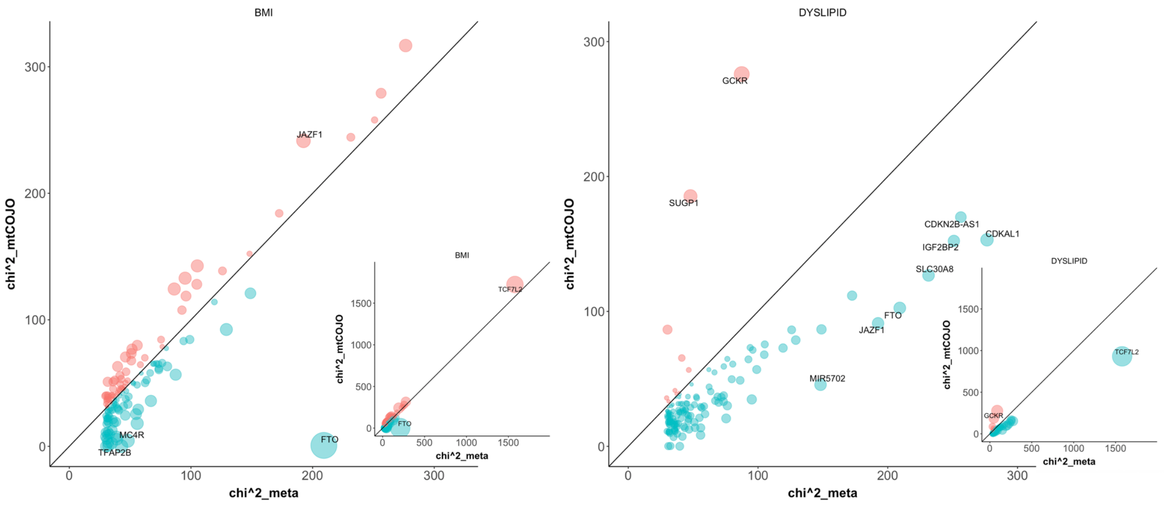


**Supplementary Figure 14** Test-statistics at the 139 T2D loci conditioning on BMI or Dyslipidaemia by a GCTA-mtCOJO analysis vs. those from the original meta-analysis. Each circle represents a locus. Shown on the x-axis are the chi-squared statistics from the original meta-analysis and those on the y-axis are the chi-square statistics from the mtCOJO analysis. The size of a circle reflects the difference in chi-square statistic between meta-analysis and mtCOJO analysis. The loci with relatively large differences are labelled with the names of their nearest genes.

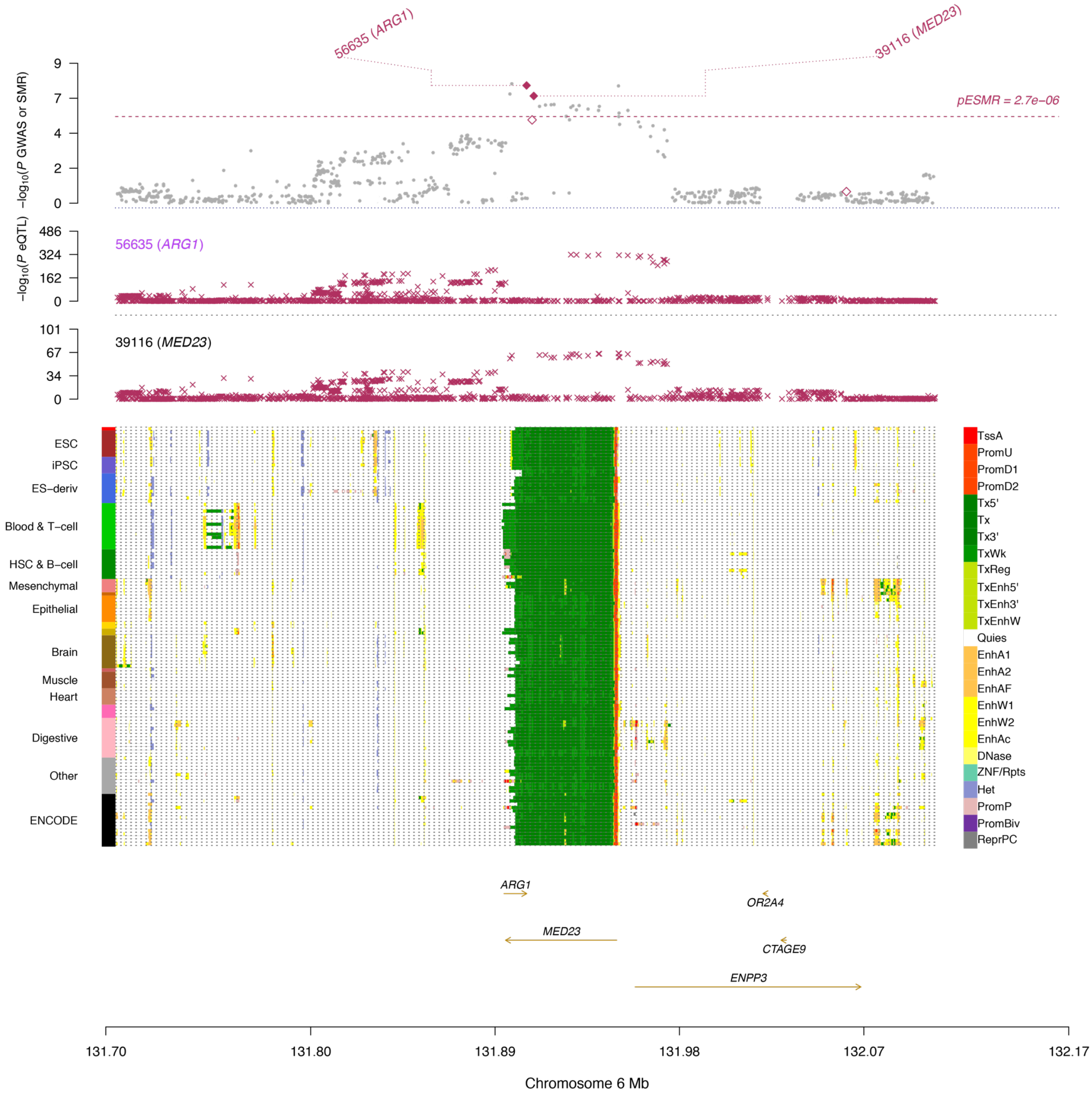


**Supplementary Figure 15** Prioritizing genes and regulatory elements at the *ARG1* locus. Shown are the results from the SMR analysis that integrates data from GWA and eQTL studies. The top plot shows -log10(*P*-value) of SNPs from the GWAS meta-analysis for T2D. Red diamonds represent -log10(*P*-value) from the SMR tests for associations of gene expression probes with T2D. Solid diamonds represent the probes not rejected by the HEIDI test. The second plot shows -log10(*P*-value) of the SNP association for gene expression probe 56635 (tagging *ARG1*) and 39166 (tagging *MED23*). The bottom plot shows 25 chromatin state annotations (indicated by colours) of 127 samples from Roadmap Epigenomics Mapping Consortium (REMC) for different primary cells and tissue types (rows).
